# Temporally controlled nervous system-to-gut signaling bidirectionally regulates longevity in *C. elegans*

**DOI:** 10.1101/2024.04.12.589179

**Authors:** Lingxiu Xu, Chengxuan Han, Lei Chun, X.Z. Shawn Xu, Jianfeng Liu

## Abstract

The nervous system modulates aging by secreting signaling molecules to cell-nonautonomously regulate the physiological state of distal tissues such as the gut. However, the underlying mechanisms are not well understood. Here, using *C. elegans* as a model, we identified two distinct neuroendocrine signaling circuits through which motor neurons signal the gut in early life to shorten lifespan but in mid-late life to extend lifespan. Both circuits employ the same neurotransmitter acetylcholine (ACh), while recruiting two different gut ACh receptors ACR-6 and GAR-3 to regulate the transcription factor DAF-16 and HSF-1 in early and mid-late life, respectively. Strikingly, the gut expression of ACR-6 is restricted to early life, whereas that of GAR-3 is confined to mid-late life, providing a potential mechanism for the temporal control of the two circuits. These results identify a novel mechanism that empowers the nervous system to bidirectionally regulate longevity by differentially signaling the gut at different life stages.

## Introduction

Aging is a complex physiological process characterized by a progressive functional decline in tissues and organs, leading to the onset of age-related diseases and probability of death^1,2^. The aging process occurs body wide in multiple tissues/organs in a coordinated manner, though different tissues/organs may age at different rates^3-8^. Recent work has uncovered an increasingly important role for the nervous system in orchestrating such a coordinated aging process through tissue-tissue communications^5,9-14^. The nervous system mediates tissue-tissue communications by secreting various signaling molecules such as small neurotransmitters and neuropeptides, which modulate the physiological state of distal tissues. For example, in *C. elegans*, the nervous system can detect and process internal stress signals from mitochondria and external sensory cues, such as temperature and odors, and then communicates with the distal tissue gut to regulate aging in a cell-nonautonomous manner^14-21^. How the nervous system signals distal tissues to regulate aging remains a question under intensive investigation.

The nervous system modulates aging in a complex fashion. Research in the past decade has uncovered both negative and positive impact of the nervous system on aging. The notion that the output of neural cells, including both neurons and glial cells, plays an important role in aging has gained consensus^19,22,23^, though conflicting results have been reported as to whether enhanced neural output is beneficial or detrimental to longevity. For example, several studies have demonstrated that neuronal hyper-excitability shortens lifespan in organisms ranging from *C. elegans* to humans^24-26^. Specifically, the transcription factor REST suppresses neural gene expression in humans and mice, and overexpression of the *C. elegans* REST orthologue *spr-4* suppresses neuronal output and extends lifespan^24^. On the other hand, several studies support the opposing view that enhanced neuronal output extends lifespan. For example, genetic and pharmacological potentiation of motor neuron activity and synaptic transmission at neuromuscular junctions (NMJs) slow down motor aging and extend lifespan in *C. elegans*^27-29^. These seemingly conflicting results reflect the complexity of the role of the nervous system in aging modulation. One possibility is that the nervous system can both negatively and positively regulate aging, depending on the context; however, whether and how the nervous system does so is not well understood.

In this study, we sought to address these questions in *C. elegans*, a widely used genetic model organism for the study of aging. Using the *C. elegans* motor nervous system as a model, we developed a strategy to temporally manipulate the motor neuron output at different life stages. We identified two distinct neuroendocrine signaling circuits through which neuronal output inhibits longevity in early life but promotes it in mid-late life. Both circuits require the neurotransmitter acetylcholine (ACh) as a signaling molecule that signals the gut via two different gut ACh receptors, ACR-6 and GAR-3, to regulate two different transcription factors, DAF-16 and HSF-1, at early and mid-late life stages, leading to shortened and extended lifespan, respectively. Strikingly, the expression of the ACh receptors ACR-6 and GAR-3 in the gut undergoes a temporal switch in early and mid-late life, providing a potential mechanism to temporally control the activity of the two circuits. Our results define the molecular and circuit mechanisms by which the nervous system bidirectionally regulates longevity by differentially signaling the gut at different life stages, highlighting an exquisite role of the nervous system in aging modulation.

## Results

### Cholinergic motor neurons differentially regulate lifespan at different stages of worm life

To ascertain whether the nervous system promotes or inhibits longevity, we focused on the motor nervous system using *C. elegans* as a model. Previous work has uncovered a role of *C. elegans* cholinergic but not GABAergic motor neurons in the ventral nerve cord in lifespan regulation^28-30^. To further explore the role of cholinergic motor neurons in lifespan control, we ablated the output of these motor neurons by expressing tetanus toxin (TeTx) as a transgene in these neurons using the *acr-2* promoter. TeTx cleaves the SNARE protein synaptobrevin, thereby blocking the release of signaling molecules such as neurotransmitters and neuropeptides from the neurons of interest^31^.

Overexpression of TeTx induces characteristic phenotypes of cholinergic deficiency, such as developmental delay and severe locomotion impairment^32^, yet does not compromise muscle function (Figure S1A). Surprisingly, such intervention led to a complex effect on the population survival curve by reducing both early mortality and the proportion of long-lived individuals (Figure 1A). Specifically, the 25% lifespan of these worms was prolonged, while their 75% and maximal lifespan were slightly shortened, leading to a mean lifespan slightly increased or unchanged compared to that of wild-type worms. This suggests that inhibiting cholinergic motor neurons may exert temporally distinct effects on survival, leading to decreased individual variation in lifespan. To test this model, we performed the converse experiment by promoting the output of cholinergic motor neurons. This was achieved by expressing in these neurons a transgene encoding syntaxin(T254I), a gain-of-function form of *Drosophila* syntaxin, another SNARE protein required for neuronal exocytosis^33^. Expression of this syntaxin mutant enhances synaptic release without causing vesicle depletion, thereby promoting neuronal output^33^, an approach that has previously been successfully applied to manipulate neuronal output in lifespan assays^20^. This transgene simply shortened lifespan without causing a pleotropic effect (Figure 1B), and critically, without inducing motor neuron degeneration (Figure S1B). This suggests that enhancing the output from cholinergic motor neurons in early life may cause an irreversible detrimental effect that overwhelms its beneficial effect in mid-late life.

**Figure 1.**
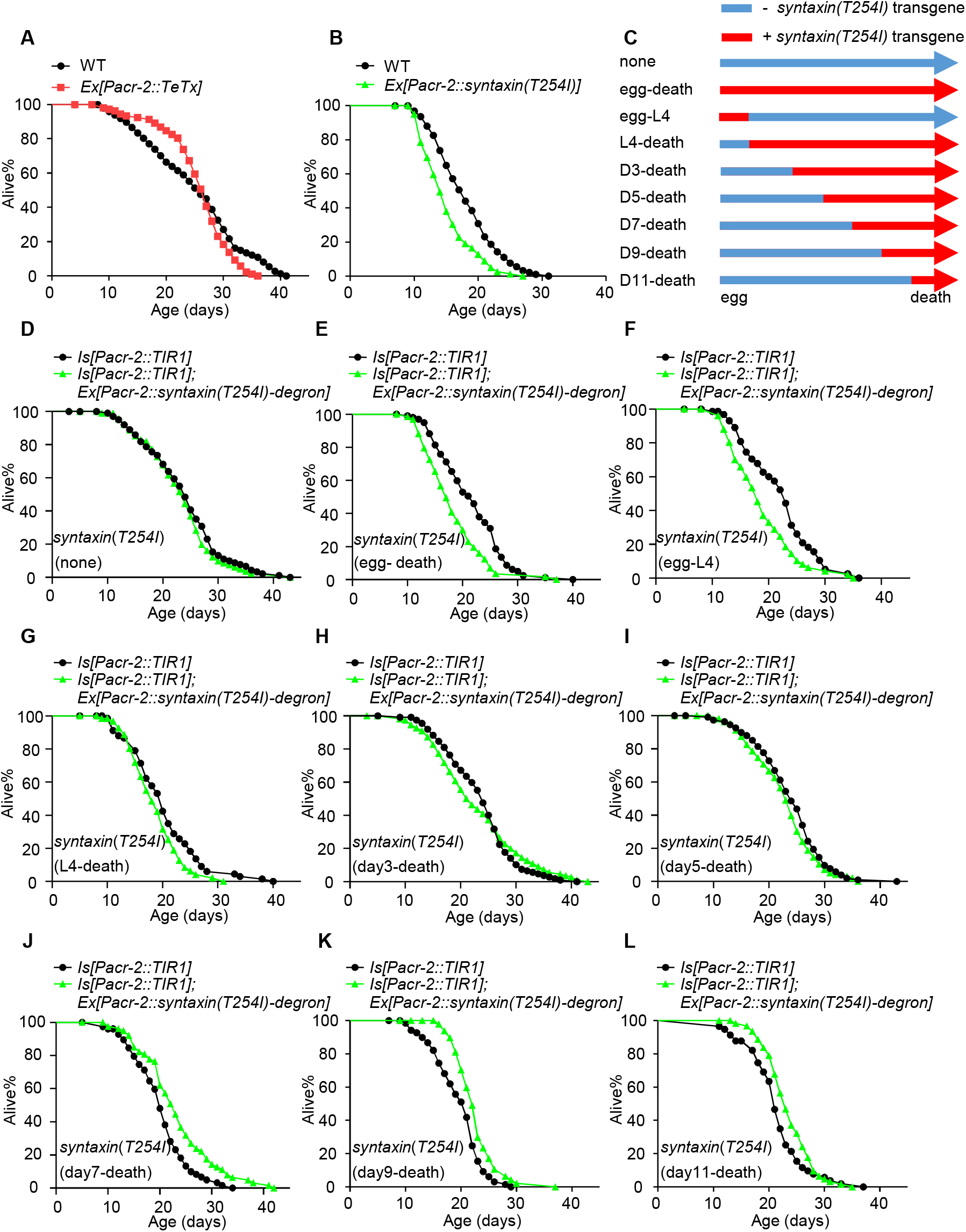
Cholinergic motor neurons differentially regulate lifespan at different stages of worm life. (A) Ablating the output of cholinergic motor neurons promotes survival at early life stage, whereas inhibits survival at later stages. Tetanus toxin (TeTx) was expressed as a transgene in cholinergic motor neurons using *acr-2* promoter to block exocytosis from these neurons. (B) Promoting the output of cholinergic motor neurons shortens lifespan. The gain-of-function form of *Drosophila* syntaxin(T254I) was expressed as a transgene in cholinergic motor neurons using *acr-2* promotor to potentiate exocytosis from these neurons. (C) Schematic describing the assay. The gain-of-function form of *Drosophila* syntaxin(T254I) fused with a degron tag was expressed as a transgene in cholinergic motor neurons to promote exocytosis from these neurons. Transgenic worms were then crossed with a line stably expressing TIR1 in these neurons. Expression of Syntaxin(T254I) was suppressed by 0.5 mM auxin treatment and induced by subsequently removing auxin. (D) *Syntaxin(T254I)* transgene has slight effect on lifespan when suppressing its expression throughout the entire worm life. (E) *Syntaxin(T254I)* transgene shortens lifespan when inducing its expression throughout the entire worm life. (F) *Syntaxin(T254I)* transgene shortens lifespan when inducing its expression from egg to L4 larval stage. (G-I) *Syntaxin(T254I)* transgene modestly affects lifespan when inducing its expression from L4 larval stage (G), Day 3 (H) or Day 5 (I) adulthood. (J-K) *Syntaxin(T254I)* transgene extends lifespan when inducing its expression from Day 7 (J), Day 9 (K) or Day 11 (L) adulthood. See Table S1 for detailed statistical analysis of lifespan data. Logrank (Kaplan-Meier) was used to calculate p values.

To further characterize the temporal effect of cholinergic motor neurons on lifespan, we sought to vary the output of cholinergic motor neurons in a temporally controlled manner. To do so, we employed an auxin-induced protein degradation (AID) system to express syntaxin(T254I) fused with a degron tag as a transgene in cholinergic motor neurons^34^. Transgenic worms were then crossed with a line stably expressing TIR1 in these neurons. TIR1 is the plant ubiquitin ligase, an essential component of the AID system^34^. Expression of syntaxin(T254I) can be suppressed by auxin treatment and induced by subsequently removing auxin. Comparable outcomes were obtained with both 0.1 mM and 0.5 mM auxin treatments (Figure S1C-1D). In doing so, we were able to potentiate the output of cholinergic motor neurons in a temporally controlled manner by expressing syntaxin(T254I) at different stages of worm life (Figure 1C). As a control, we first shut down the expression of *syntaxin(T254I)* transgene throughout the entire worm life by treating worms with auxin at all life stages, and found that this had no notable effect on lifespan (Figure 1D). Conversely, enhancing the output of cholinergic neurons throughout the entire worm life (no auxin treatment) shortened lifespan, consistent with the view that the detrimental effect resulting from enhanced motor neuron output in early life overwrites its beneficial effect at later stages (Figure 1E). Indeed, enhancing the output of cholinergic motor neurons during the larval stage only (no auxin from egg to L4) shortened lifespan, confirming the detrimental effect of motor neuron output on lifespan in early life (Figure 1F). Notably, this detrimental effect began to fade when motor neuron output was instead enhanced during the adult stage (auxin from egg to adult stage), and eventually worms became long-lived when motor neuron output was enhanced after Day 5 adulthood (Figure 1G-1L). These data reveal a dichotomous role of cholinergic motor neurons in lifespan regulation, shortening adult lifespan in early life but extending it in mid-late life.

### ACh mediates the lifespan-shortening effect of cholinergic motor neurons in early life

How do cholinergic motor neurons differentially control lifespan at different life stages? First, we focused on their lifespan-shortening effect. Cholinergic motor neurons release both small neurotransmitters and neuropeptides^32,35-37^. To determine which type of signaling molecules might be released from motor neurons to shorten lifespan in early life, we first tested *unc-31* mutant worms, which are defective in secreting neuropeptides^38-40^. Loss of *unc-31* exhibited no defect on the ability of cholinergic motor neurons to shorten lifespan, indicating that neuropeptides may not play an important role in this process (Figure 2A and 2B). This led us to test small neurotransmitters. Cholinergic motor neurons release ACh, and indeed, loss of *unc-17* gene, which encodes the sole *C. elegans* vesicular ACh transporter^41^, eliminated the lifespan-shortening effect of these motor neurons (Figure 2A and 2C). In *unc-17* mutant worms, no ACh is expected to be released from cholinergic motor neurons. As a control, we also tested mutants deficient in other types of small neurotransmitters, including glutamate (*eat-4*), GABA (*unc-25*), serotonin (*tph-1*), dopamine (*cat-2*), tyramine (*tdc-1*), and octopamine (*tbh-1*), but detected no effect, with the exception of *tph-1*, which showed a modest, partial suppression of the phenotype (Figure S2A-S2F). This observation suggests that the lifespan effects of cholinergic signaling can be modulated by serotonin. These results uncover a key role of the neurotransmitter ACh in mediating the lifespan-shortening effect of cholinergic motor neurons in early life.

**Figure 2.**
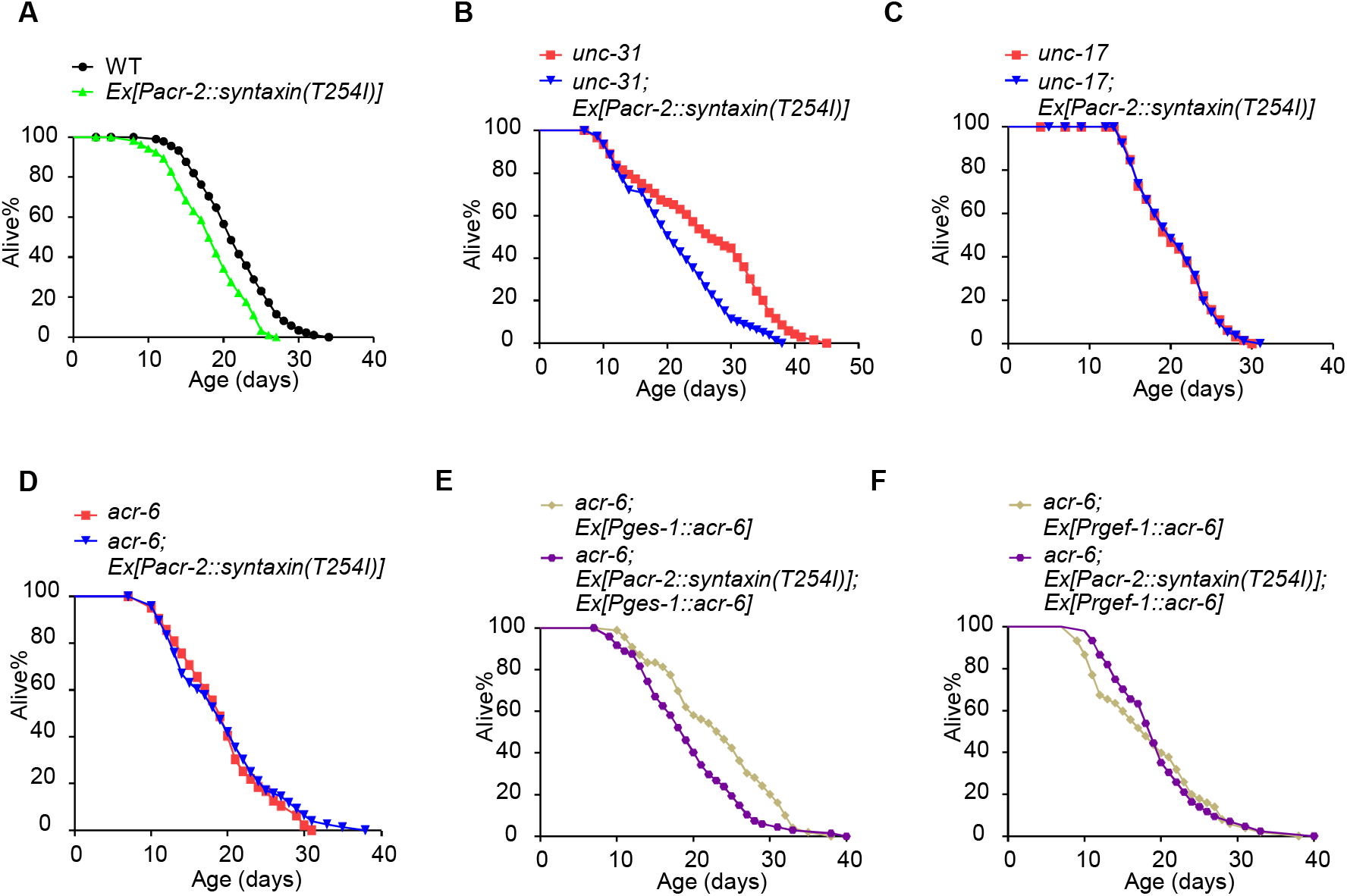
ACh and the nicotinic AChR ACR-6 mediate the lifespan-shortening effect of cholinergic motor neurons in early life. (A) *Syntaxin(T254I)* transgene shortens lifespan. (B) Loss of *unc-31* exhibits no defect on the ability of *syntaxin(T254I)* transgene to shorten lifespan. (C) Loss of *unc-17* blocks the ability of *syntaxin(T254I)* transgene to shorten lifespan. (D-F) Loss of *acr-6* blocks the ability of *syntaxin(T254I)* transgene to shorten lifespan (D), a phenotype rescued by transgenic expression of wild-type *acr-6* gene in the intestine (E), but not in neurons (F). The *ges-1* and *rgef-1* promoters were used to drive *acr-6* gene expression in the intestine and neurons, respectively. See Table S1 for detailed statistical analysis of lifespan data. Logrank (Kaplan-Meier) was used to calculate p values.

### The nicotinic AChR ACR-6 acts in the intestine to mediate the lifespan-shortening effect of cholinergic motor neurons

ACh is expected to act through ACh receptors (AChRs). To identify the AChRs that mediate the lifespan-shortening effect of cholinergic motor neurons, we examined all of the 37 *C. elegans* AChR genes, including nicotinic AChRs (nAChRs), muscarinic AChRs (mAChRs), and ACh-gated Cl^-^ channels^32^. Notably, RNAi of several ACh receptors such as *acr-11* appears to shorten wild-type lifespan, whereas RNAi of several other ACh receptors such as *acr-9* extends wild-type lifespan, suggesting lifespan-modulating potential of ACh receptors (Figure S3). By screening strains with each of these 37 AchRs knocked down by RNAi, we identified three candidate genes: *acr-6* (nAChR), *acr-8* (nAChR) and *gar-2* (mAChR) (Figure S3). To validate these candidates, we examined their mutants. Mutation in *acr-6* blocked the ability of cholinergic motor neurons to shorten lifespan, while mutations in the other two candidates did not (Figure 2D and Figure S4A-S4F). This identifies a key role for the nAChR ACR-6 in mediating the lifespan-shortening effect of cholinergic motor neurons. Previous data has shown that cholinergic signaling and ACR-6 may control pharyngeal pumping^42^. As expected, we found that *acr-6* mutation slightly reduced pumping rates (Figure S4G).

We then asked in which tissue ACR-6 regulates lifespan. Transgenic expression of wild-type *acr-6* gene in the intestine (Figure 2E), but not in neurons (Figure 2F), rescued the *acr-6* mutant phenotype, indicating that ACR-6 acts in the intestine. This suggests that ACh released from cholinergic motor neurons may shorten lifespan by acting on its cognate receptor ACR-6 in the intestine, revealing a nervous system-to-gut signaling axis.

### Cholinergic motor neurons require the transcription factor DAF-16 in the intestine to regulate lifespan

Aging pathways tend to converge on a subset of transcription factors^12^. We went on to search for a transcription factor that functions in the intestine to mediate the lifespan-shortening effect of cholinergic motor neurons in early life. RNAi of *daf-16*, which encodes a FOXO transcription factor that is a key regulator of lifespan, did not further shorten the lifespan of worms expressing *syntaxin(T254I)* transgene (Figure 3A and 3B); as a control, RNAi of other transcription factor such as *hsf-1*/HSF1, *skn-1*/Nrf2 *and pha-4*/FOXA did (Figure 3C-3E). This suggests that *daf-16* may act in the same pathway as cholinergic motor neurons. DAF-16 regulates lifespan by controlling the expression of its target genes. As expected, the expression level of *sod-3* and *mtl-1*, two commonly characterized DAF-16 target genes, was upregulated in transgenic worms deficient in releasing ACh from cholinergic motor neurons (Figure 3F), and downregulated in transgenic worms with enhanced ACh release from cholinergic motor neurons (Figure S5A), consistent with the notion that DAF-16 acts downstream of cholinergic motor neurons. To obtain further evidence, we assessed the subcellular localization pattern of DAF-16::GFP fusion and found that *acr-6* RNAi notably promoted nuclear translocation of DAF-16, confirming that ACh signaling inhibits DAF-16 activity (Figure S5B).

**Figure 3.**
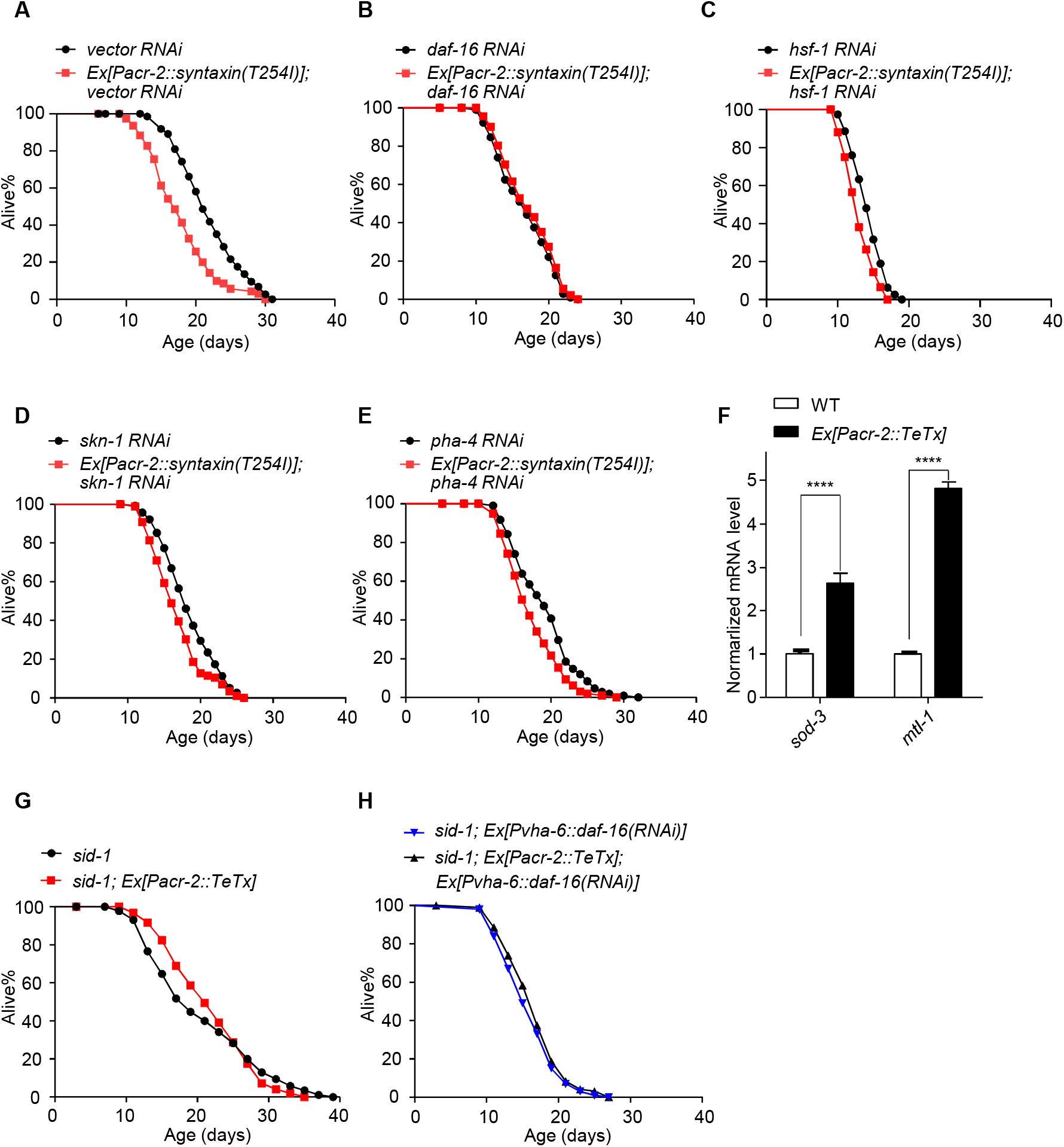
The FOXO transcription factor DAF-16 is required in the intestine for cholinergic motor neurons to regulate lifespan in early life. (A) *Syntaxin(T254I)* transgene shortens lifespan. (B-E) RNAi of *daf-16* does not further shorten the lifespan of *syntaxin(T254I)* transgenic worms, while RNAi of *hsf-1* (C), *skn-1* (D) and *pha-4* (E) does. (F) qPCR analysis of DAF-16 target genes. qPCR reactions were run in triplicates for each genotype. Day 1 adult worms were detected. Each experiment was repeated three times. Error bars represent s.e.m. ****p< 0.0001 (ANOVA with Bonferroni’s test). (G-H) RNAi of *daf-16* in the intestine (H) abolishes the ability of TeTx transgene to regulate lifespan (G). dsRNA against *daf-16* was expressed as a transgene specifically in the intestine using *vha-6* promoter. These experiments were carried out in a *sid-1* mutant background where systemic effect of RNAi is absent. See Table S1 for detailed statistical analysis of lifespan data. Log-rank (Kaplan-Meier) was used to calculate p values.

DAF-16 is known to regulate lifespan primarily in the intestine^4^. To determine whether DAF-16 also functions in the intestine, we performed tissue specific RNAi experiments in *sid-1* mutant background to restrict RNAi to specific tissues. dsRNA against *daf-16* was expressed as a transgene specifically in the intestine. RNAi of *daf-16* in the intestine abolished the ability of cholinergic motor neurons to regulate lifespan (Figure 3G, 3H and Figure S5C-5E). Thus, similar to ACR-6, DAF-16 also appears to function in the intestine, suggesting that the intestine is an important target tissue to which cholinergic motor neurons signals to regulate lifespan. It also suggests that cholinergic motor neurons shorten lifespan by regulating DAF-16 activity in the intestine.

Collectively, the data presented above suggests a model in which cholinergic motor neurons regulate lifespan through a neuroendocrine signaling circuit (Figure 6D, below). In this circuit, cholinergic motor neurons signal the intestine via the neurotransmitter ACh that acts through its cognate receptor ACR-6 to regulate DAF-16 in the intestine, leading to a shortened lifespan (Figure 6D). Because this lifespan-shortening effect results from enhanced motor neuron output in early life and overwrites its beneficial effect at later stages, we propose this signaling circuit mediates the lifespan-shortening effect in early life.

### ACh mediates the lifespan-extending effect of cholinergic motor neurons in mid-late life

Having characterized the mechanisms by which cholinergic motor neurons signal the intestine to shorten lifespan in early life, we then set out to explore how these motor neurons extend lifespan in mid-late life. Given that ACh is the only neurotransmitter known to be released by these motor neurons and its role in mediating motor neuron-to-gut signaling in early life, we asked whether ACh also plays a role in mediating the lifespan-extending effect of these motor neurons in mid-late life. Again, we resorted to the AID system to specifically enhance the output of cholinergic motor neurons in mid-late life by temporally restricting the expression of *syntaxin(T254I)* transgene at this stage. We found that cholinergic motor neurons lost the ability to extend lifespan at mid-late life stage in *unc-17* mutant worms that are deficient in ACh release, revealing an essential role of ACh in mediating the lifespan-extending effect of these motor neurons in mid-late life and suggesting it as a key signaling molecule in this process (Figure 4A and 4B). Thus, the same neurotransmitter ACh appears to mediate both the lifespan-shortening and extending effects of cholinergic motor neurons.

**Figure 4.**
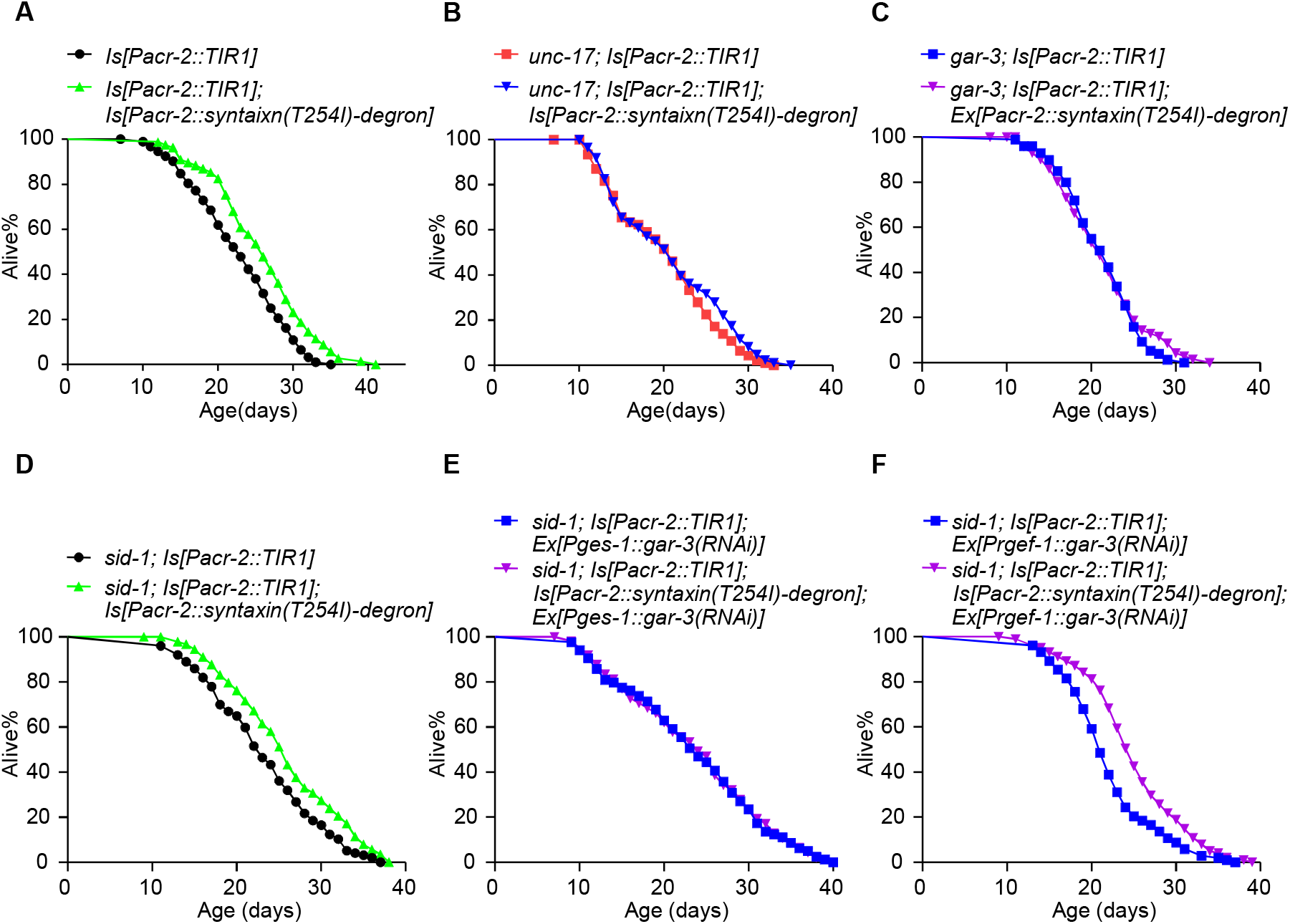
ACh and the intestinal muscarinic AChR GAR-3 mediate the lifespan-extending effect of cholinergic motor neurons in mid-late life. (A) Enhancing the output of cholinergic motor neurons in mid-late life extends lifespan. The *syntaxin(T254I)* transgene was induced from Day 7 adulthood as described in Figure 1C. (B-C) Loss of *unc-17* (B) or *gar-3* (C) blocks the ability of cholinergic motor neurons to extend lifespan in mid-late life. (D-F) RNAi of *gar-3* in the intestine (E), but not in neurons (F), abolishes the ability of cholinergic motor neurons to extend lifespan in mid-late life (D). dsRNA against *gar-3* was expressed as a transgene specifically in the intestine and neurons using *ges-1* and *rgef-1* promoter, respectively. These experiments were carried out in a *sid-1* mutant background where systemic effect of RNAi is absent. See Table S1 for detailed statistical analysis of lifespan data. Log-rank (Kaplan-Meier) was used to calculate p values.

### The muscarinic AChR GAR-3 acts in the intestine to mediate the lifespan-extending effect of cholinergic motor neurons in mid-late life

We then asked how the same neurotransmitter ACh secreted from the same type of neurons mediates two opposite effects on lifespan. To address this question, we re-screened all the 37 AChR genes (Figure S6). Two candidate genes emerged from the screen: *gar-2* and *gar-3*, which encode two different mAChRs. RNAi of *gar-2* or *gar-3* completely suppressed the ability of cholinergic motor neurons to extend lifespan in mid-late life (Figure S7A-S7C). *gar-2* and *gar-3* mutant worms exhibited a similar phenotype (Figure S7D, S7E and Figure 4C), suggesting that GAR-2 and GAR-3 may be the receptors for ACh that mediate the lifespan-extending effect of cholinergic motor neurons.

To determine in which tissue GAR-2 and GAR-3 act to regulate lifespan, we specifically knocked down *gar-2* and *gar-3* genes in the intestine and neurons, as both genes are known to be expressed in these two tissues^43-46^. RNAi of *gar-3* in the intestine (Figure 4D and 4E), but not in neurons or the muscle (Figure 4D-4F, and Figure S8A, S8D-S8E), abolished the ability of cholinergic motor neurons to extend lifespan at mid-late life stage. Thus, GAR-3 may function in the intestine to regulate lifespan. Surprisingly, RNAi of *gar-2* in the muscle (Figure S8A-S8C), but not in neurons or the intestine (Figure S7F-S7H) had an effect on the ability of cholinergic motor neurons to extend lifespan in mid-late life, indicating that GAR-2 acts in the muscle to regulate lifespan. Given our focus on neuron-gut signaling, we decided to focus on GAR-3 for further characterizations. These results identify GAR-3 as a candidate AChR that acts in the intestine to mediate the effect of the neurotransmitter ACh and cholinergic motor neurons on lifespan extension in mid-late life.

### Cholinergic motor neurons require the transcription factor HSF-1 in the intestine to regulate lifespan in mid-late life

In light of our observation that the transcription factor DAF-16 acts in the intestine to regulate lifespan in early life, we wondered if DAF-16 also regulates lifespan in mid-late life. Surprisingly, RNAi of *daf-16* and other transcription factors such as *skn-1* and *pha-4* failed to suppress the lifespan-extending effect of cholinergic motor neurons (Figure S9). By contrast, RNAi of *hsf-1* completely abolished the ability of these motor neurons to extend lifespan in mid-late life (Figure 5A and 5B), suggesting that HSF-1, rather than DAF-16, is required by cholinergic motor neurons to regulate lifespan at mid-late life stage. Additional support came from the observation that the expression of *hsp-16*.*2* and *pat-10*, two commonly characterized HSF-1 target genes, were upregulated in long-lived worms in which the output of cholinergic motor neurons was enhanced at mid-late life stage (Figure 5C). HSF-1 is known to regulate lifespan primarily in the intestine and neurons^47,48^. To identify in which tissue HSF-1 functions, we inactivated *hsf-1* specifically in the intestine or neurons by RNAi (Figure 5D-5F). RNAi of *hsf-1* in the intestine but not in neurons abolished the ability of cholinergic motor neurons to extend lifespan at mid-late life stage. Thus, HSF-1 appears to function in the intestine to regulate lifespan, suggesting that cholinergic motor neurons extend lifespan in mid-late life through HSF-1 in the intestine.

**Figure 5.**
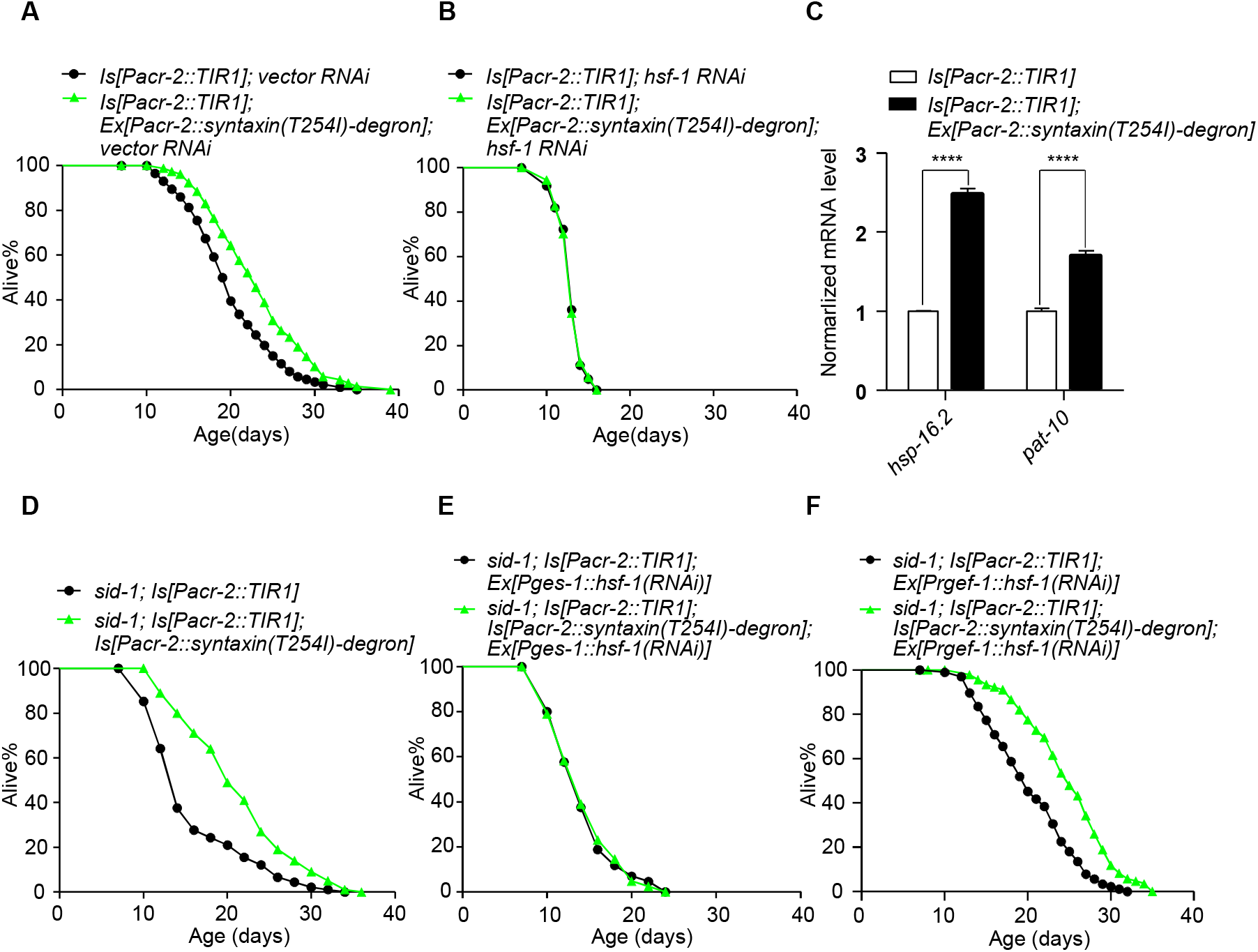
The transcription factor HSF-1 is required in the intestine for cholinergic motor neurons to regulate lifespan in mid-late life. (A-B) Enhancing the output of cholinergic motor neurons in mid-late life extends lifespan (A). This effect was fully suppressed by RNAi of *hsf-1* (B). The *syntaxin(T254I)* transgene was induced from Day 7 adulthood as described in Figure 1C. (C) qPCR analysis of HSF-1 target genes. qPCR reactions were run in triplicates for each genotype. Day 10 adult worms were detected. Each experiment was repeated three times. Error bars represent s.e.m. ****p< 0.0001 (ANOVA with Bonferroni’s test). (D-F) RNAi of *hsf-1* in the intestine (E), but not in neurons (F), abolishes the ability of cholinergic motor neurons to extend lifespan in mid-late life (D). dsRNA against *hsf-1* was expressed as a transgene specifically in the intestine and neurons using *ges-1* and *rgef-1* promoter, respectively. These experiments were carried out in a *sid-1* mutant background where systemic effect of RNAi is absent. See Table S1 for detailed statistical analysis of lifespan data. Logrank (Kaplan-Meier) was used to calculate p values.

In summary, this set of data suggests a model in which cholinergic motor neurons signal the distal tissue intestine in mid-late life to extend lifespan through a distinct neuroendocrine circuit (Figure 6D). Similar to the lifespan-shortening circuit described above, this circuit is also formed by cholinergic motor neurons and the intestine, and employs the same neurotransmitter ACh; however, it recruits a different AChR GAR-3 to regulate a different type of transcription factor HSF-1 in the intestine, leading to the opposite outcome in lifespan (Figure 6D).

**Figure 6.**
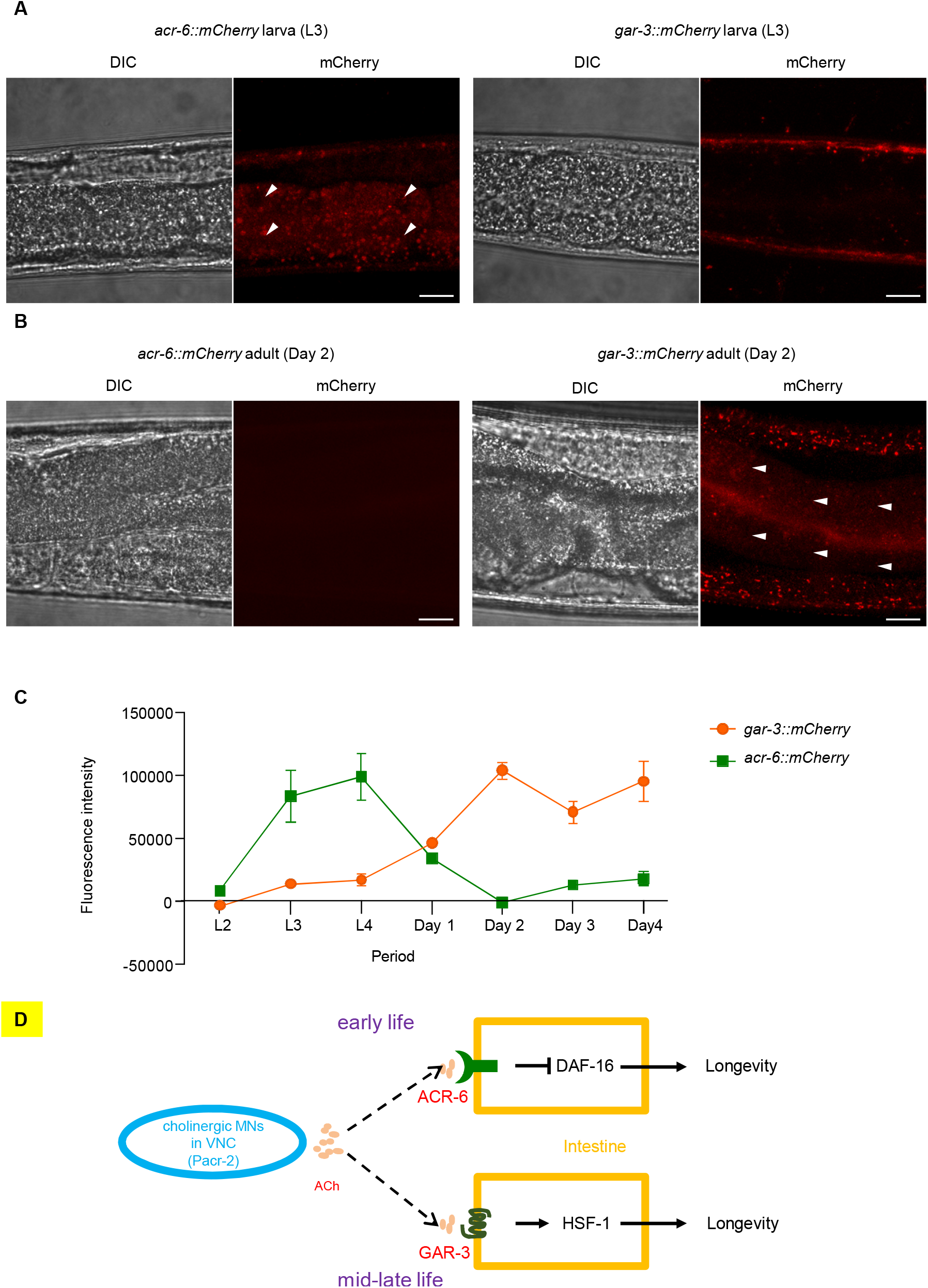
Temporal control of ACR-6 and GAR-3 expression in the intestine at different life stages. (A-B) The expression pattern of endogenous ACR-6 and GAR-3 proteins at L3 Larva stage (A) and Day 2 adult stage (B). Arrowheads point to the area of the intestine. Scale bars, 20 μm. (C) Quantification curves summarizing the data in (A) and (B). The entire intestinal area was selected for measurement. Error bars represent s.e.m. n ≥ 10. (D) A schematic model illustrating the two cholinergic motor neurons-to-gut signaling circuits that act in early life to shorten lifespan and in mid-late life to extend lifespan.

### Temporal control of ACR-6 and GAR-3 expression in the intestine at different life stages

The identification of two distinct neuroendocrine circuits provides a potential explanation to the opposing effects of cholinergic motor neurons on lifespan at different life stages. However, it prompts an important question as to how these two circuits are temporally controlled at different stages of worm life. One possibility is that the expression of some components in the two circuits are temporally regulated at different life stages. To test this idea, we sought to examine the expression of the two AChRs ACR-6 and GAR-3 in the intestine. Using CRISPR/Cas9 genome editing, we inserted a mCherry tag at the C-terminal end of the *acr-6* and *gar-3* gene to label the endogenous ACR-6 and GAR-3 protein. At the larval stage, ACR-6::mCherry fluorescence was detected in the intestine, while GAR-3::mCherry was not (Figure 6A). Remarkably, beginning at the adult stage, ACR-6::mCherry expression started to decrease and became undetectable at Day 2 adulthood (Figure 6B). On the other hand, GAR-3::mCherry began to show expression at the adult stage and became prominent at Day 2-4 adulthood (Figure 6A and 6B). We were unable to examine GAR-3::mCherry expression at Day 5 adulthood and beyond, due to the interference by increasingly bright gut auto-fluorescence even in the *glo-4(ok623)* background with reduced gut auto-fluorescence^49^ (Figure S10). Nevertheless, these results unveil a clear temporal switch in the intestinal expression of ACR-6 and GAR-3 at different life stages (Figure 6C). Though additional mechanisms may also contribute, such a dichotomy in the intestinal expression of ACR-6 and GAR-3 at larval and adult stages provides a potential mechanism that underlies the temporal control of the two neuroendocrine circuits, leading two opposing outcomes in lifespan.

## Discussion

Research in the past decade has revealed an increasingly important role of the nervous system in orchestrating a body-wide aging process by signaling distal tissues, particularly the gut^3,4,11,12,14-19,30^. How the nervous system signals the gut to regulate aging, however, is not well understood, and work in this area has yielded some seemingly inconsistent data. This points to a model in which nervous system-gut signaling can both positively and negatively regulate aging, highlighting a complex role of the nervous system-gut signaling axis in aging^17,20,30^. Nervous system-gut communications may differentially regulate aging through spatial mechanisms. For example, different neurons may play distinct roles in aging by positively and/or negatively regulating longevity via secreting distinct neurotransmitters and neuropeptides^17,18,20,50-52^. By sharp contrast, whether and how the nervous system temporally regulates aging is poorly understood. Here, using the *C. elegans* motor nervous system as a model, we identified a temporal mechanism by which the nervous system both positively and negatively regulates lifespan by temporally controlling nervous system-to-gut signaling at different life stages. Specifically, we identified two distinct neuroendocrine signaling circuits by which cholinergic motor neurons act to shorten lifespan in early life, whereas extending lifespan in mid-late life. The two circuits employ the same neurotransmitter ACh, but recruit two different AChRs ACR-6 and GAR-3 that act in the intestine to regulate two different transcription factors DAF-16 and HSF-1 to shorten and extend lifespan at early and mid-late life stage, respectively. To the best of our knowledge, this represents the first such example illustrating how the nervous system may differentially regulate lifespan by temporally controlling nervous system-to-gut signaling. Nevertheless, it is still unclear whether other neuronal populations share similar temporal regulatory mechanisms. A previous study reported that inhibiting cholinergic neurons activity (using *unc-17* promoter) extends lifespan regardless of timing^24^, which is different to the temporal lifespan regulation we observed in cholinergic motor neurons (using *acr-2* promoter). This discrepancy is likely due to differences in subsets of neurons, as the *unc-17* promoter labels a broad repertoire of cholinergic neurons, while the *acr-2* promoter mainly marks cholinergic motor neurons^53^. Thus, the distinct lifespan-modulating effects of cholinergic motor neurons may be overshadowed by opposing contributions from other cholinergic subtypes when a mixed population is manipulated. Alternatively, both activation and inhibition of cholinergic activity may perturb neurotransmission balance, leading to similar effects on lifespan^54^. It will be interesting to test these hypotheses in future studies.

One interesting observation is that the expression level of ACR-6 and GAR-3 in the intestine features a temporal switch at the larval and adult stages. The appearance followed by disappearance of ACR-6 expression in the intestine at the larval and adult stages nicely correlate with the temporal window under which cholinergic motor neurons shorten lifespan. This provides a potential mechanism by which cholinergic motor neurons shorten lifespan in early life. Remarkably, the temporal expression pattern of GAR-3 in the intestine follows a reciprocal pattern: it is not expressed at the larval stage and only appears at the adult stage, providing a potential mechanism for the lifespan-extending effect exerted by cholinergic motor neurons in mid-late life. It should be pointed out that the intestinal expression of GAR-3 emerges at Day 1 adulthood, earlier than the time point at which the beneficial role of cholinergic motor neurons began to take effect. Several factors might have contributed to this delay. For example, the lifespan-shortening signaling in the intestine might not have been completely shut off during early adulthood, particularly considering that the role of the transcription factor DAF-16 might be long-lasting. This might antagonize the effect of the lifespan-extending signaling initiated by GAR-3 in early adulthood. Additionally, though GAR-3 expression can be detected in early adulthood, it might take some time for it to fully engage other signaling components, which might cause a delay for the lifespan-extending signaling to take effect. Future studies are needed to elucidate the mechanism that directs the temporal switch of ACR-6 and GAR-3 expression in the intestine.

Though the temporally-controlled dichotomous expression pattern of ACR-6 and GAR-3 in the intestine provides a potential mechanism underlying the opposing effects of cholinergic motor neurons on longevity, we do not exclude the possibility that additional mechanisms may apply. For example, the expression pattern of other components in the two neuroendocrine circuits may undergo a similar temporal shift; similarly, their function may also be subjected to temporal control. An alternative mechanism based on differential levels of cholinergic signaling could also contribute to the observed lifespan effects. Our work provides an entry point to understand how temporal mechanisms may contribute to the complex roles of the nervous system as well as nervous system-gut communications in aging.

## Acknowledgments

We thank Wenxin Jia, Jiyong Meng, Yang Fu, Jinghan Shi and Guanghui Hu for technical assistance. Some strains were obtained from the CGC. This work was supported by the NSFC (32171146 to L.C. and 31130028, 31225011 and 31420103909 to J.L.) and funds from the University of Michigan.

## Author contributions

L.X., C.H. and L.C. conducted the experiments and analyzed the data. L.X., C.H., L.C., X.Z.S.X., and J.L. interpreted the data. L.C., X.Z.S.Z., and J.L. wrote the article with assistance from other authors.

## Data availability

The datasets generated and analyzed during this study are either included within the manuscript or are available from the authors upon request.

## MATERIALS AND METHODS

### Strains and genetics

Wild-type: N2. *Ex263a[Pacr-2::TeTx::sl2::yfp]. Is4a[Pacr-2::TIR1::sl2::yfp]. Is4a; Ex329a[Pacr-2::syntaxin(T254I)-degron::sl2::mCherry]. Ex294a[Pacr-2::syntaxin(T254I)::sl2::mCherry]. unc-31(e169). unc-31; Ex294a. cat-2(e1112). cat-2; Ex294a. tbh-1(n3247). tbh-1; Ex294a. tph-1(mg280). tph-1; Ex294a. eat-4(ky5). eat-4; Ex294a. unc-25(e156). unc-25; Ex294a. tdc-1(n3419). tdc-1; Ex294a. unc-17(e245). unc-17; Ex294a. acr-6(ok3117). acr-6; Ex294a. acr-8(ok1240). acr-8; Ex294a. gar-2(ok520). gar-2; Ex294a. acr-6; Ex321a[Pges-1::acr-6::sl2::yfp]. acr-6; Ex294a; Ex321a. acr-6; Ex324a[Prgef-1::acr-6::sl2::yfp]. acr-6; Ex294a; Ex324a. sid-1(qt9). sid-1; Ex263a. sid-1; Ex380a[Pvha-6::daf-16 (RNAi)+Pvha-6::sl2::mCherry]. sid-1; Ex263a; Ex380a. sid-1; Ex371a[Prgef-1::daf-16 (RNAi)+Prgef-1::sl2::mCherry]. sid-1; Ex263a; Ex371a. Is4a; Is5a[Pacr-2::syntaxin(T254I)-degron::sl2::mCherry]. unc-17; Is4a. unc-17; Is4a; Is5a. gar-3(gk337); Is4a. gar-3; Is4a; Ex329a. sid-1; Is4a. sid-1; Is4a; Is5a. sid-1; Is4a; Ex359a[Pges-1::gar-3(RNAi)+Pges-1::sl2::cfp]. sid-1; Is4a; Is5a; Ex364a[Pges-1::gar-3(RNAi)+Pges-1::sl2::cfp]. sid-1; Is4a; Ex395a[Prgef-1::gar-3(RNAi)+Prgef-1::sl2::cfp]. sid-1; Is4a; Is5a; Ex398a[Prgef-1::gar-3(RNAi)+Prgef-1::sl2::cfp]. gar-2; Is4a. gar-2; Is4a; Ex329a. sid-1; Is4a; Ex357a[Pges-1::gar-2(RNAi)+Pges-1::sl2::cfp]. sid-1; Is4a; Is5a; Ex363a[Pges-1::gar-2(RNAi)+Pges-1::sl2::cfp]. sid-1; Is4a; Ex390a[Prgef-1::gar-2(RNAi)+Prgef-1::sl2::cfp]. sid-1; Is4a; Is5a; Ex392a[Prgef-1::gar-2(RNAi)+Prgef-1::sl2::cfp]. sid-1; Is4a; Ex373a[Pges-1:hsf-1(RNAi)+Pges-1::sl2:cfp]. sid-1; Is4a; Is5a; Ex374a[Pges-1:hsf-1(RNAi)+Pges-1::sl2:cfp]. sid-1; Is4a; Ex384a[Prgef-1::hsf-1(RNAi)+Prgef-1::sl2::cfp]. sid-1; Is4a; Is5a; Ex385a[Prgef-1::hsf-1(RNAi)+Prgef-1::sl2::cfp]. acr-6(xu129a[acr-6::mCherry]). gar-3(xu130a[gar-3::mCherry]). ieSi57[Peft-3::TIR1]. ieSi57[Peft-3::TIR1]; Ex329a. Is4a; Ex847a[Pvha-6::daf-16 (RNAi)+Pvha-6::sl2::mCherry] line 1. Is4a; Is5a; Ex847a[Pvha-6::daf-16 (RNAi)+Pvha-6::sl2::mCherry] line 1. Is4a; Ex848a[Pvha-6::daf-16 (RNAi)+Pvha-6::sl2::mCherry] line 2. Is4a; Is5a; Ex848a[Pvha-6::daf-16 (RNAi)+Pvha-6::sl2::mCherry] line 2. Is4a; Ex859a[Pmyo-3::gar-2(RNAi)+Pmyo-3::sl2::mCherry] line 1. Is4a; Is5a; Ex859a[Pmyo-3::gar-2(RNAi)+Pmyo-3::sl2::mCherry] line 1. Is4a; Ex860a[Pmyo-3::gar-2(RNAi)+Pmyo-3::sl2::mCherry] line 2. Is4a; Is5a; Ex860a[Pmyo-3::gar-2(RNAi)+Pmyo-3::sl2::mCherry] line 2. Is4a; Ex857a[Pmyo-3::gar-3(RNAi)+Pmyo-3::sl2::mCherry] line 1. Is4a; Is5a; Ex857a[Pmyo-3::gar-3(RNAi)+Pmyo-3::sl2::mCherry] line 1. Is4a; Ex858a[Pmyo-3::gar-3(RNAi)+Pmyo-3::sl2::mCherry] line 2. Is4a; Is5a; Ex858a[Pmyo-3::gar-3(RNAi)+Pmyo-3::sl2::mCherry] line 2. zIs356[daf-16::gfp + roller]*.

Mutant strains were outcrossed at least four times before use. The *acr-6, acr-8, acr-15, acr-16, acr-19, acr-21, acr-24, gar-1, acc-1, acc-2, acc-4, deg-3, lev-8, pbo-5, eat-2* and *unc-63* RNA interference (RNAi) clones were generated in the laboratory, while other RNAi clones were from the Ahringer library and were confirmed by sequencing.

Microinjections were performed using standard protocols. Each plasmid DNA listed above in the transgenic line was injected at a concentration of 50 ng/μL. Each marker for RNAi was co-injected at a concentration of 25 ng/μL.

### Worm synchronization

Strains were grown at 20°C for at least three generations before lifespan determination. Twenty Day 2 adult worms were transferred to fresh 60 mm nematode growth medium (NGM) plates. The adult worms were removed from the plates after laying eggs for 4 hours. The plates were placed at 20°C for two days. L4 larva were used to start the normal lifespan or RNAi assays.

### Lifespan assay

Lifespan assays were performed on 60 mm NGM plates at 20°C as previously described^15^. ∼120 worms of each genotype were used for lifespan assays and transferred every other day to fresh NGM plates. Survival rate was scored every 1 – 2 days. Worms were censored if they crawled off the plate, bagged, or exhibited protruding vulva. In all cases, the first day of adulthood was scored as day 1. Lifespan data was analyzed with GraphPad Prism 8 (GraphPad Software, Inc.) and IBM SPSS Statistics 21 (IBM, Inc.). Log-rank (Kaplan-Meier) was used to calculate *P* values. All lifespan assays were repeated at least twice. 2 μg/mL 5-Fluoro-2’-deoxyuridine (FUDR) was included in assays involving TeTx transgene worms, *unc-31* and *unc-17* mutant worms, which show a defect in egg laying.

### RNA interference (RNAi) assay

RNAi was performed using the RNAi-compatible OP50 bacterial strain OP50(xu363) and HT115 as previously described^55^. RNAi plates included carbenicillin (25 μg/mL) and isopropyl-β-D-thiogalactoside (0.5 mM). OP50(xu363) or HT115 bacteria with vector or RNAi plasmid were seeded on RNAi plates 2 days before experiment.

### Auxin treatment

Auxin treatment was performed by transferring worms to bacteria-seeded NGM plates containing 0.5 mM natural auxin indole-3-acetic acid (G&K Scientific) as previously described^34^. Auxin was dissolved in cholesterol solution (25 g/L) and KP buffer (2.5 g/L) to avoid additional ethanol. Cholesterol solution and KP buffer with auxin were added to NGM agar that has cooled down to about 60°C before pouring the plates. Expression of syntaxin(T254I) can be suppressed by auxin treatment and restored in 24 hours following auxin removal.

### Microscopy

Different stage worms were anesthetized with 5 mM sodium azide (Sigma) on 2% agarose pads. Confocal images were acquired by FLUOVIEW FV3000 series of confocal laser scanning microscopes (Olympus) equipped with a 100X objective as Z-stacks and were processed using ImageJ (NIH). Maximum intensity Z-projection images were shown to illustrate the endogenous ACR-6 and GAR-3 protein expression pattern. To quantify the ACR-6 and GAR-3 protein expression level in the intestine, maximum intensity Z-projection were used and the entire intestine area were selected by ImageJ.

### CRISPR genome editing

ACR-6::mCherry and GAR-3::mCherry are both knockin alleles made by injecting RNP mixtures as described previously^56^. In brief, repair templates were amplified using unmodified primers with 70 bp homology arms against the target genes from linker-mCherry plasmid. The RNP mixture was assembled at 37°C for 15 min and contained Cas9 protein, tracrRNA, and crRNA (all from IDT). crRNA corresponding DNA target sequences are 5’-ATTCACGGATTGATAGCAAT-3’ (*acr-6*), and 5’-ATTCACGGATTGATAGCAAT-3’ (*gar-3*). The double-strand donor templates were melted right before adding them to the assembled RNPs ^57^. Microinjection quality was scored using a co-injection marker, PRF4::*rol-6 (su1006)* plasmid. In general, 50 P0 were injected, singled, and maintained at 25°C for 3 days. Two plates with the highest number of F1 rollers (usually more than 20 rollers for a good injection) were selected, and 48 F1 rollers from each plate were picked and singled for further genotyping^56^. The positive insertions were confirmed by Sanger sequencing. Using this CRISPR-RNP technology, the coding sequence of mCherry and a 27 bp linker sequence were inserted immediately after the last amino acid codon of the *acr-6* gene on chromosome I and of *gar-3* gene on chromosome V, respectively.

### Levamisole paralysis assay

Worms were synchronized at the L4 larval stage one day prior to the assay. The following day, 20-30 adult worms were transferred to a plate with 1 mM levamisole hydrochloride and prodded every 10 minutes for 2 hours to assess movement. Worms that showed no response to this harsh touch were classified as paralyzed.

### Nuclear translocation of DAF-16::GFP

L3 larva stage worms were anesthetized with 5 mM sodium azide (Sigma) on 2% agarose pads. Confocal images were acquired by FLUOVIEW FV3000 series of confocal laser scanning microscopes (Olympus) equipped with a 20X objective as Z-stacks. Image processing and quantification of DAF-16::GFP nuclear localization were carried out using ImageJ (NIH). For each worm, the nuclear localization level was calculated as the percentage of intestinal cells showing nuclear GFP signal.

