## Supplemental Figure 1-10 for "Temporally controlled nervous system-to-gut signaling bidirectionally regulates longevity in *C. elegans*"

Figure S1

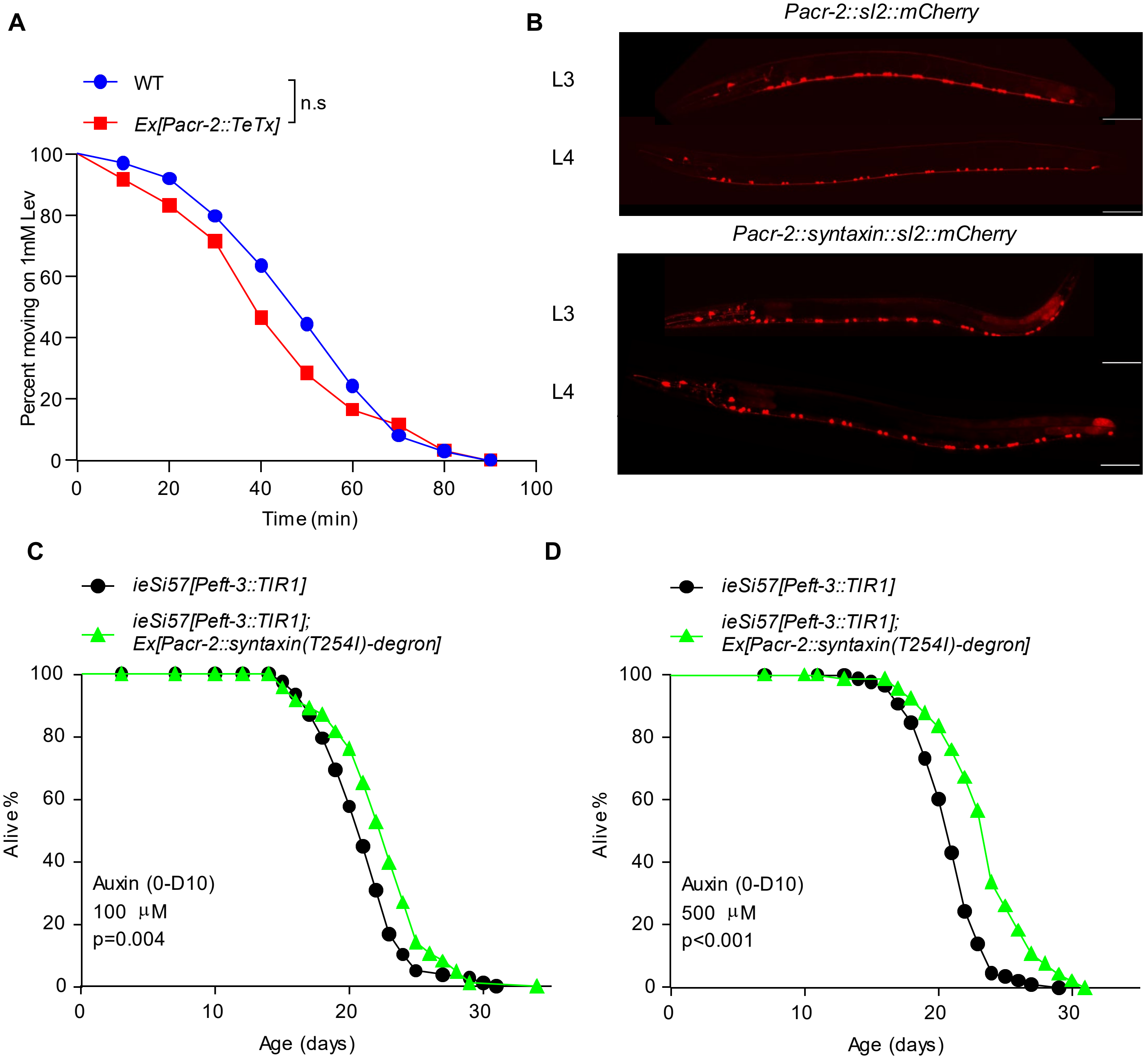


**Figure S1. Cholinergic motor neuron manipulation affects neuronal output without compromising muscle function and motor neuron morphology.**

(A) Ablating the output of cholinergic motor neurons does not affect muscle function. Tetanus toxin (TeTx) was expressed as a transgene in cholinergic motor neurons using *acr-2* promoter to block exocytosis from these neurons. 20-30 Day 1 adult worms were transferred to a plate with 1 mM levamisole hydrochloride and monitored over 2 hours by prodding every 10 minutes. prodded every 10 min over a 2 hours-period to determine if they retained the ability to move. Worms that showed no movement in response to this harsh touch were classified as paralyzed. Log-rank (Kaplan-Meier) was used to calculate p values.P=0.1662. ns: no significant difference. (B) Promoting the output of cholinergic motor neurons does not cause motor neuron degeneration. The gain-of-function form of Drosophila syntaxin(T254I) was expressed as a transgene in cholinergic motor neurons using *acr-2* promotor to potentiate exocytosis from these neurons (lower panels). For the control, the *Pacr-2::sl2::mCherry* transgene was expressed instead (upper panels).The labeled motor neuronal cell bodies were calculated at specific time points. Scale bars, 50 μm (C-D) Syntaxin(T254I) transgene extends lifespan when its expression is induced from day 10 of adulthood, following prior suppression from egg to Day 10 adulthood with 100 μM (C) or 500 μM (D) auxin. The gain-of-function form of Drosophila syntaxin(T254I) fused with a degron tag was expressed as a transgene in cholinergic motor neurons to promote exocytosis from these neurons. Transgenic worms were then crossed with a line stably expressing TIR1 in soma. Expression of Syntaxin(T254I) was suppressed by 100 μM or 500 μM auxin treatment and restored by subsequently removing auxin.

Figure S2

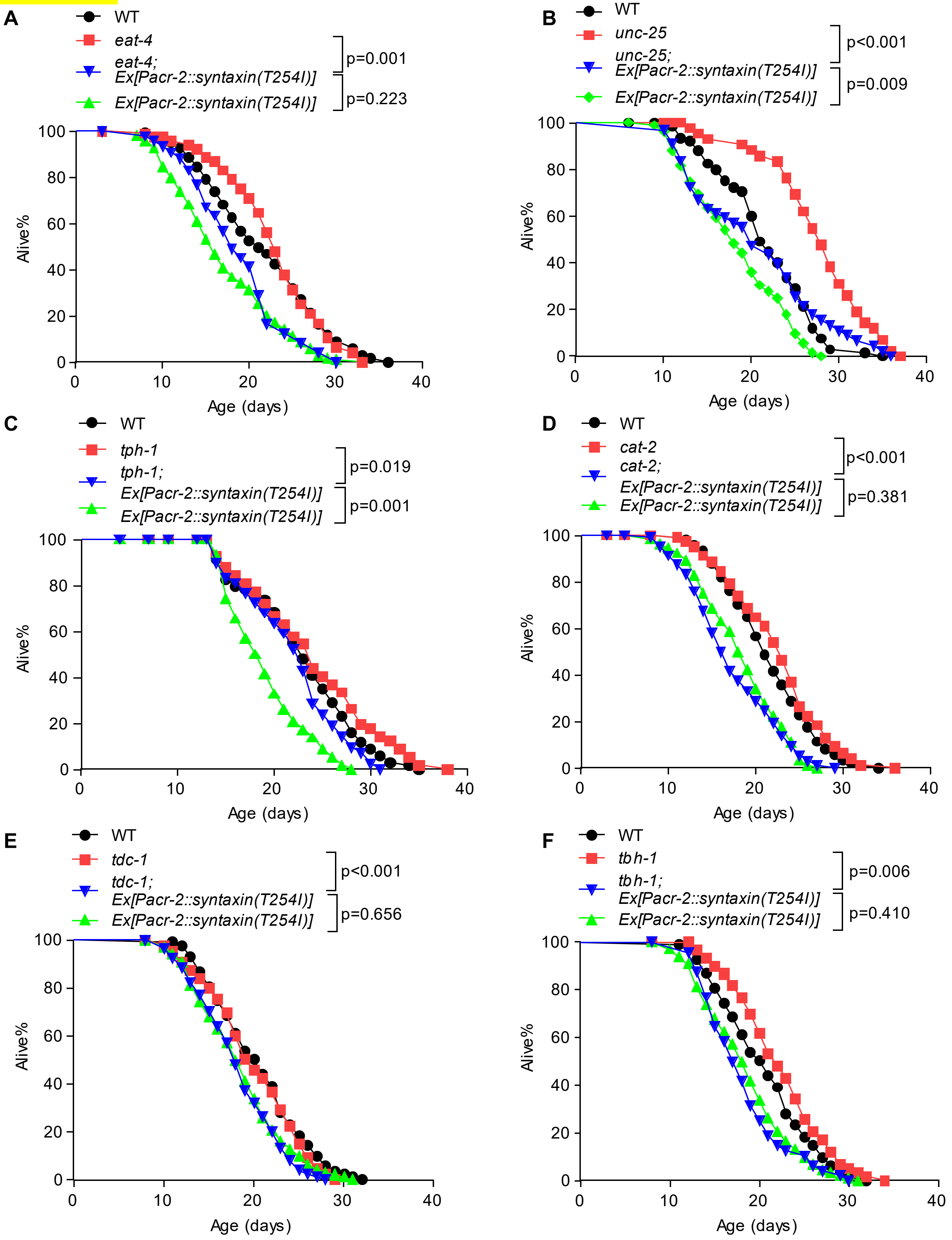


**Figure S2. Mutants deficient in small neurotransmitters other than ACh exhibit no or modest defect in the ability of cholinergic motor neurons to shorten lifespan in early life.**

**(A-F)** Loss of *eat-4* (A), *unc-25* (B), *tph-1* (C), *cat-2* (D), or *tdc-1* (E) exhibits no defect in the ability of cholinergic motor neurons to shorten lifespan, while *tbh-1* (F) showed a modest suppression of the phenotype. All lifespan assays were performed at 20 °C and were repeated twice. Log-rank (Kaplan-Meier) was used to calculate p values.

Figure S3

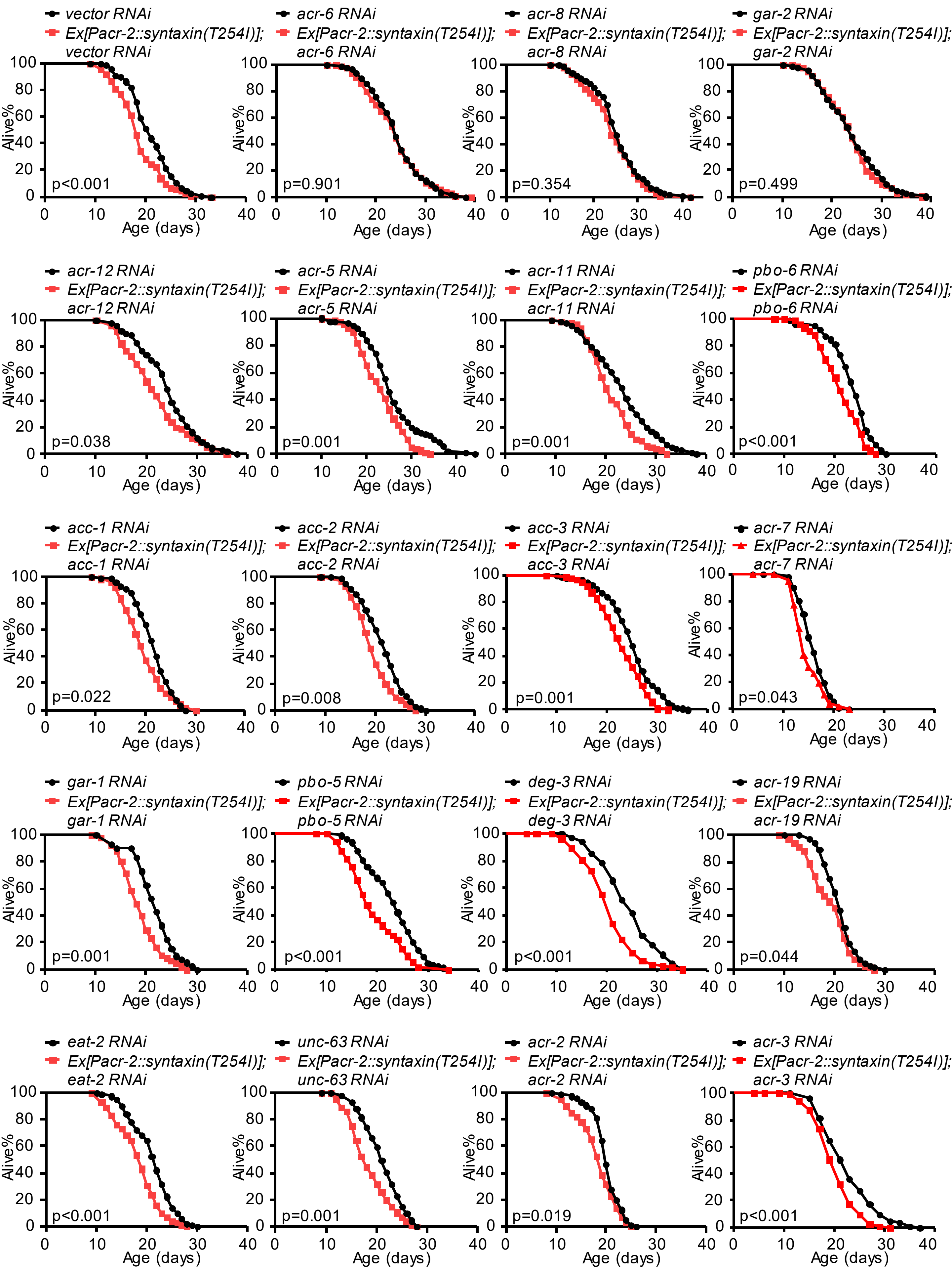
vector RNAi
Ex[Pacr-2::syntaxin(T254I)]; vector RNAi
p<0.001
acr-6 RNAi
Ex[Pacr-2::syntaxin(T254I)]; acr-6 RNAi
p=0.901
acr-8 RNAi
Ex[Pacr-2::syntaxin(T254I)]; acr-8 RNAi
p=0.354
gar-2 RNAi
Ex[Pacr-2::syntaxin(T254I)]; gar-2 RNAi
p=0.499
acr-12 RNAi
Ex[Pacr-2::syntaxin(T254I)]; acr-12 RNAi
p=0.038
acr-5 RNAi
Ex[Pacr-2::syntaxin(T254I)]; acr-5 RNAi
p=0.001
acr-11 RNAi
Ex[Pacr-2::syntaxin(T254I)]; acr-11 RNAi
p=0.001
pbo-6 RNAi
Ex[Pacr-2::syntaxin(T254I)]; pbo-6 RNAi
p<0.001
acc-1 RNAi
Ex[Pacr-2::syntaxin(T254I)]; acc-1 RNAi
p=0.022
acc-2 RNAi
Ex[Pacr-2::syntaxin(T254I)]; acc-2 RNAi
p=0.008
acc-3 RNAi
Ex[Pacr-2::syntaxin(T254I)]; acc-3 RNAi
p=0.001
acr-7 RNAi
Ex[Pacr-2::syntaxin(T254I)]; acr-7 RNAi
p=0.043
gar-1 RNAi
Ex[Pacr-2::syntaxin(T254I)]; gar-1 RNAi
p=0.001
pbo-5 RNAi
Ex[Pacr-2::syntaxin(T254I)]; pbo-5 RNAi
p<0.001
deg-3 RNAi
Ex[Pacr-2::syntaxin(T254I)]; deg-3 RNAi
p<0.001
acr-19 RNAi
Ex[Pacr-2::syntaxin(T254I)]; acr-19 RNAi
p=0.044
eat-2 RNAi
Ex[Pacr-2::syntaxin(T254I)]; eat-2 RNAi
p<0.001
unc-63 RNAi
Ex[Pacr-2::syntaxin(T254I)]; unc-63 RNAi
p=0.001
acr-2 RNAi
Ex[Pacr-2::syntaxin(T254I)]; acr-2 RNAi
p=0.019
acr-3 RNAi
Ex[Pacr-2::syntaxin(T254I)]; acr-3 RNAi
p<0.001
Alive%
Age (days)

Figure S3

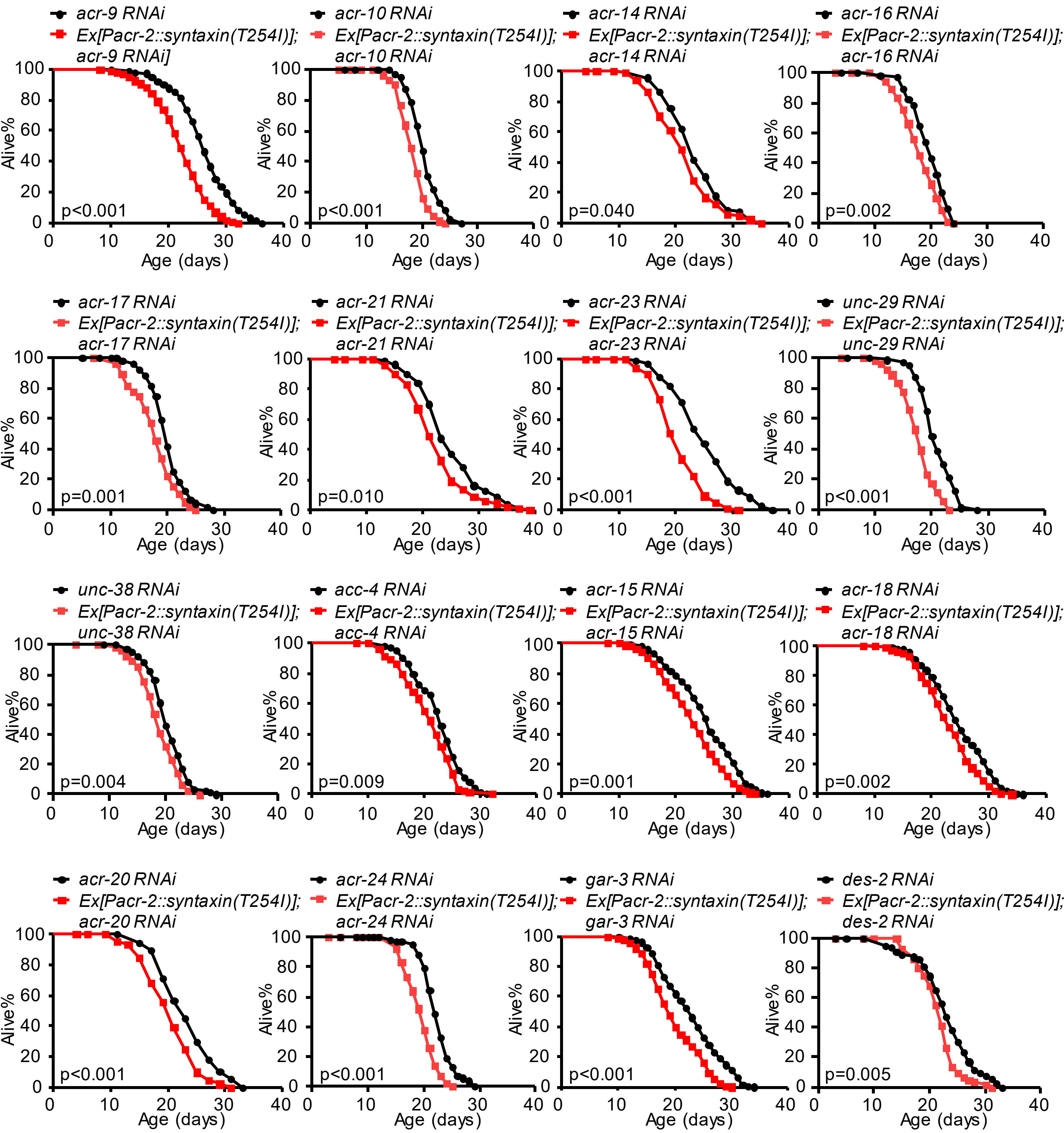


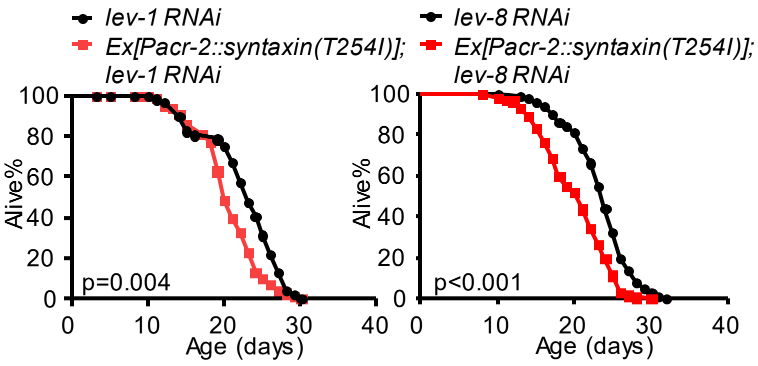


**Figure S3. RNAi screen of AChR genes that can block the lifespan-shortening effect of cholinergic motor neurons.**
For feeding RNAi experiments, NGM plates included carbenicillin (25 μg/ml) and IPTG (0.5 mM). OP50(xu363) bacteria-carrying vector or RNAi plasmid were seeded on RNAi plates 2 days before the experiment. Worms were fed RNAi bacteria from the egg stage. All lifespan assays were performed at 20 °C. Logrank (Kaplan-Meier) was used to calculate p values.

Figure S4

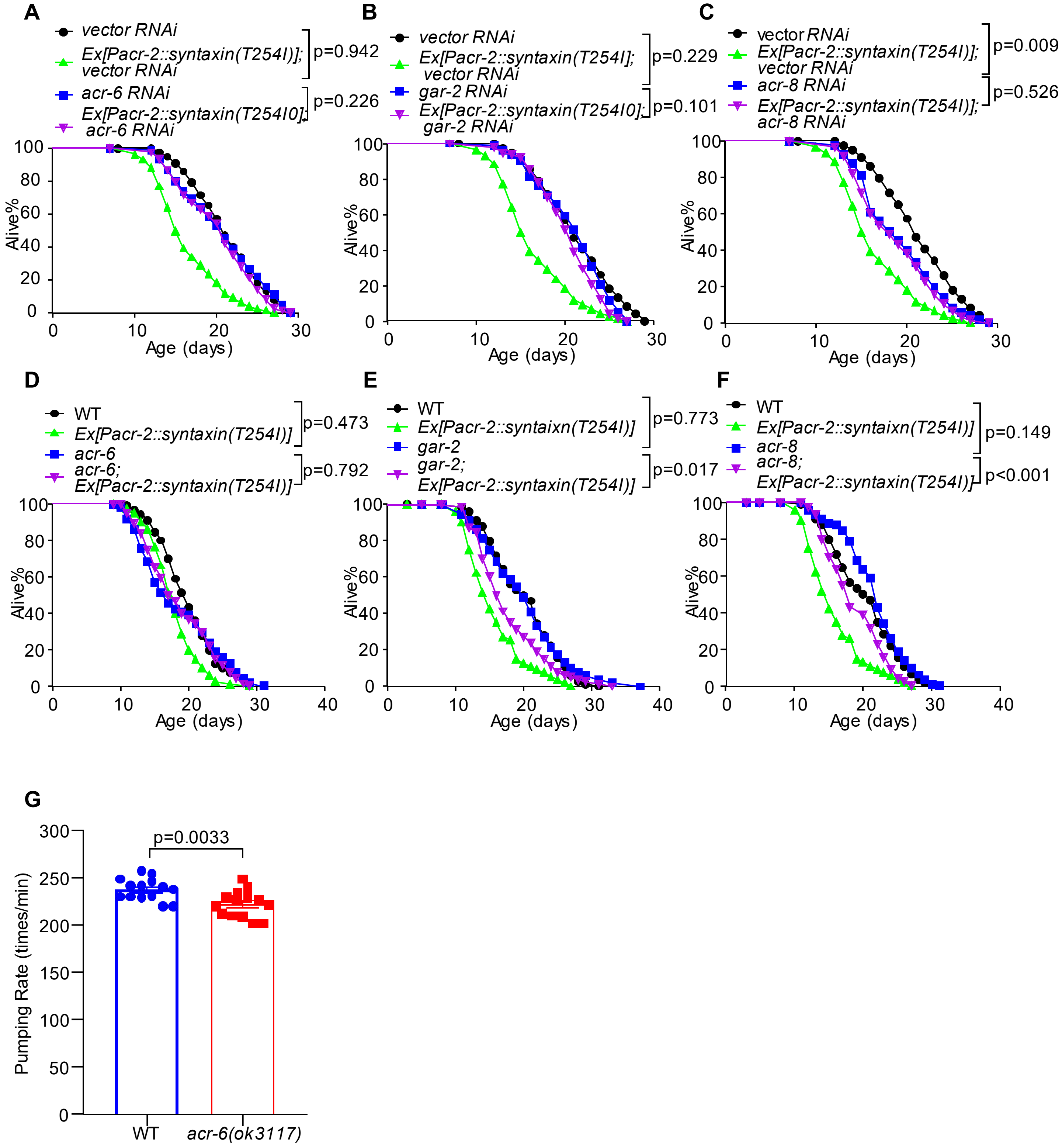


**Figure S4. Mutation in *acr-6* but not in *gar-2* or *acr-8* can block the lifespan-shortening effect of cholinergic motor neurons.**

(A-C) RNAi of *acr-6* (A), *gar-2* (B) and *acr-8* (C) genes completely suppresses the lifespan-shortening phenotype of *syntaxin(T254I)* transgene. (D-F) Mutation of *acr-6* (D) completely suppresses the lifespan-shortening phenotype of *syntaxin(T254I)* transgene, while mutations in *gar-2* (E) and *acr-8* (F) do not. For feeding RNAi experiments, NGM plates included carbenicillin (25 μg/ml) and IPTG (0.5 mM). OP50(xu363) bacteria-carrying vector or RNAi plasmid were seeded on RNAi plates 2 days before the experiment. Worms were fed RNAi bacteria from the egg stage. All lifespan assays were performed at 20 °C. Log-rank (Kaplan-Meier) was used to calculate p values. (G) Mutation of *acr-6* inhibits pumping rate. Pharyngeal pumping was measured in 20 day 1 adult worms. P=0.00333 (T-test).

Figure S5

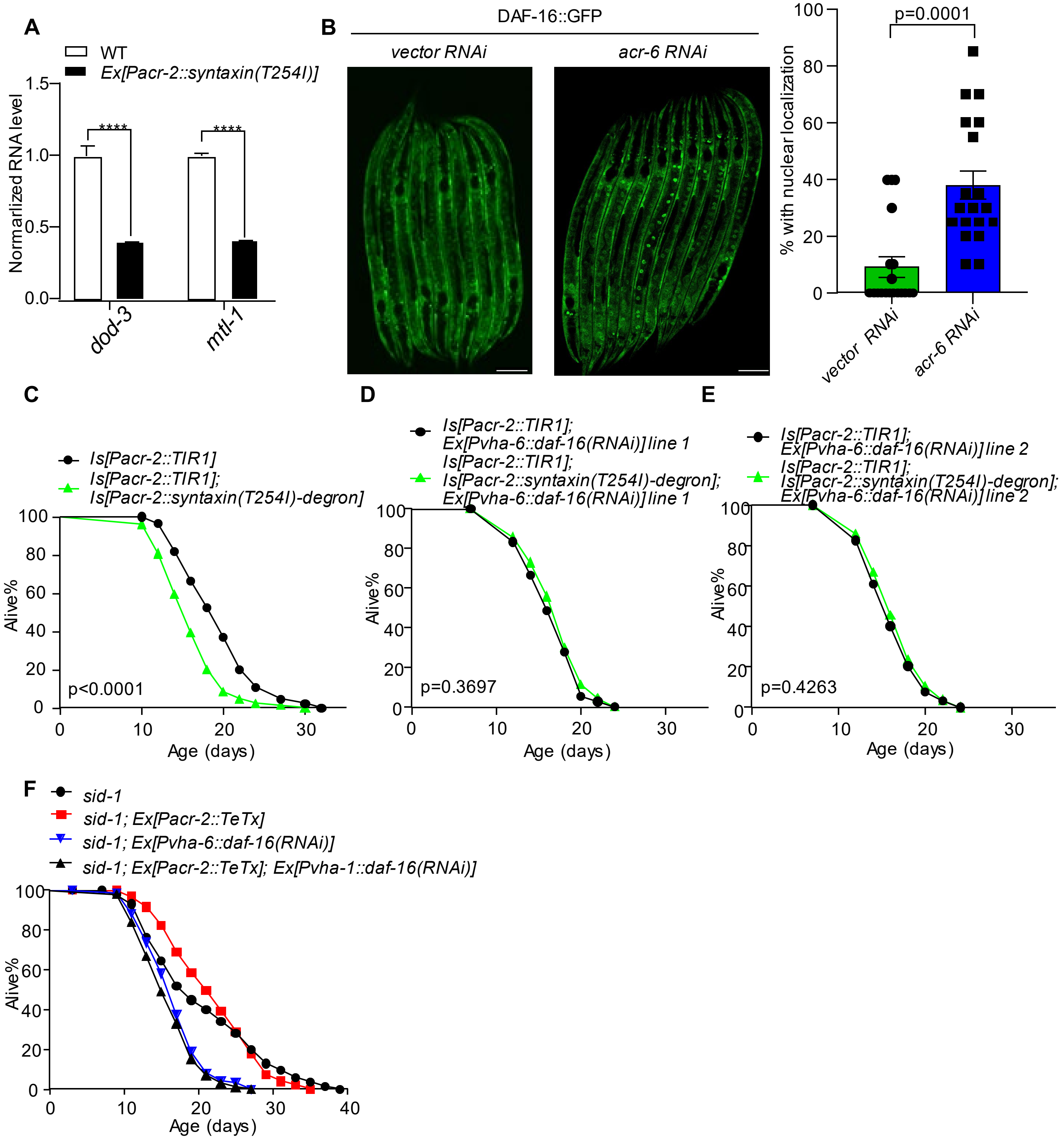


**Figure S5. The FOXO transcription factor DAF-16 is required in the intestine for cholinergic motor neurons to regulate lifespan.**

(A) qPCR analysis of DAF-16 target genes. qPCR reactions were run in triplicates for each genotype. Day 1 adult worms were detected. Each experiment was repeated three times. Error bars represent s.e.m. ****p< 0.0001 (ANOVA with Bonferroni's test). (B) Nuclear translocation of DAF-16::GFP in vector or *acr-6(RNAi)* worms. The left panels show representative images. The right panel shows the quantification of DAF-16::GFP nuclear localization, presented as the mean percentage of intestinal cells displaying nuclear GFP signal per worm. L3 larva worms were detected.Scale bars, 50 μm. p=0.0001 (t-test). (C) *Syntaxin(T254I)-degron* transgene shortens lifespan. (D-E) RNAi of *daf-16* in the intestine abolishes the ability of *Syntaxin(T254I)-degron* transgene to regulate lifespan. dsRNA against *daf-16* was expressed as a transgene specifically in the intestine using *vha-6* promoter. All lifespan assays were performed at 20 °C. Log-rank (Kaplan-Meier) was used to calculate p values. (F) The merged figure of Figure 3G and 3H.

Figure S6

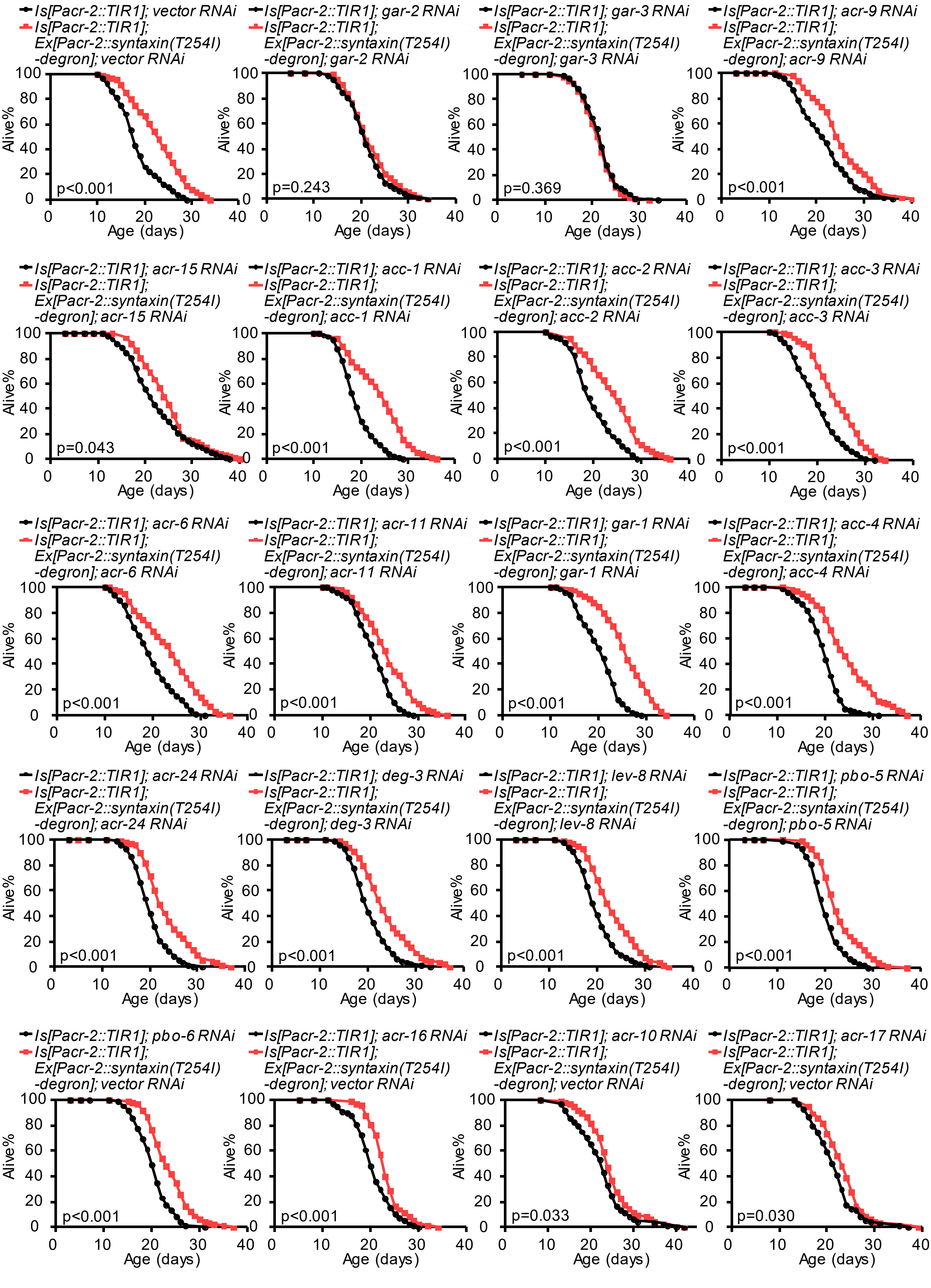

Is[Pacr-2::TIR1]; vector RNAi
Is[Pacr-2::TIR1]; Ex[Pacr-2::syntaxin(T254I) -degron]; vector RNAi
p<0.001
Is[Pacr-2::TIR1]; gar-2 RNAi
Is[Pacr-2::TIR1]; Ex[Pacr-2::syntaxin(T254I) -degron]; gar-2 RNAi
p=0.243
Is[Pacr-2::TIR1]; gar-3 RNAi
Is[Pacr-2::TIR1]; Ex[Pacr-2::syntaxin(T254I) -degron]; gar-3 RNAi
p=0.369
Is[Pacr-2::TIR1]; acr-9 RNAi
Is[Pacr-2::TIR1]; Ex[Pacr-2::syntaxin(T254I) -degron]; acr-9 RNAi
p<0.001
Is[Pacr-2::TIR1]; acr-15 RNAi
Is[Pacr-2::TIR1]; Ex[Pacr-2::syntaxin(T254I) -degron]; acr-15 RNAi
p=0.043
Is[Pacr-2::TIR1]; acc-1 RNAi
Is[Pacr-2::TIR1]; Ex[Pacr-2::syntaxin(T254I) -degron]; acc-1 RNAi
p<0.001
Is[Pacr-2::TIR1]; acc-2 RNAi
Is[Pacr-2::TIR1]; Ex[Pacr-2::syntaxin(T254I) -degron]; acc-2 RNAi
p<0.001
Is[Pacr-2::TIR1]; acc-3 RNAi
Is[Pacr-2::TIR1]; Ex[Pacr-2::syntaxin(T254I) -degron]; acc-3 RNAi
p<0.001
Is[Pacr-2::TIR1]; acr-6 RNAi
Is[Pacr-2::TIR1]; Ex[Pacr-2::syntaxin(T254I) -degron]; acr-6 RNAi
p<0.001
Is[Pacr-2::TIR1]; acr-11 RNAi
Is[Pacr-2::TIR1]; Ex[Pacr-2::syntaxin(T254I) -degron]; acr-11 RNAi
p<0.001
Is[Pacr-2::TIR1]; gar-1 RNAi
Is[Pacr-2::TIR1]; Ex[Pacr-2::syntaxin(T254I) -degron]; gar-1 RNAi
p<0.001
Is[Pacr-2::TIR1]; acc-4 RNAi
Is[Pacr-2::TIR1]; Ex[Pacr-2::syntaxin(T254I) -degron]; acc-4 RNAi
p<0.001
Is[Pacr-2::TIR1]; acr-24 RNAi
Is[Pacr-2::TIR1]; Ex[Pacr-2::syntaxin(T254I) -degron]; acr-24 RNAi
p<0.001
Is[Pacr-2::TIR1]; deg-3 RNAi
Is[Pacr-2::TIR1]; Ex[Pacr-2::syntaxin(T254I) -degron]; deg-3 RNAi
p<0.001
Is[Pacr-2::TIR1]; lev-8 RNAi
Is[Pacr-2::TIR1]; Ex[Pacr-2::syntaxin(T254I) -degron]; lev-8 RNAi
p<0.001
Is[Pacr-2::TIR1]; pbo-5 RNAi
Is[Pacr-2::TIR1]; Ex[Pacr-2::syntaxin(T254I) -degron]; pbo-5 RNAi
p<0.001
Is[Pacr-2::TIR1]; pbo-6 RNAi
Is[Pacr-2::TIR1]; Ex[Pacr-2::syntaxin(T254I) -degron]; vector RNAi
p<0.001
Is[Pacr-2::TIR1]; acr-16 RNAi
Is[Pacr-2::TIR1]; Ex[Pacr-2::syntaxin(T254I) -degron]; vector RNAi
p<0.001
Is[Pacr-2::TIR1]; acr-10 RNAi
Is[Pacr-2::TIR1]; Ex[Pacr-2::syntaxin(T254I) -degron]; vector RNAi
p=0.033
Is[Pacr-2::TIR1]; acr-17 RNAi
Is[Pacr-2::TIR1]; Ex[Pacr-2::syntaxin(T254I) -degron]; vector RNAi
p=0.030
Alive%
Age (days)

Figure S6

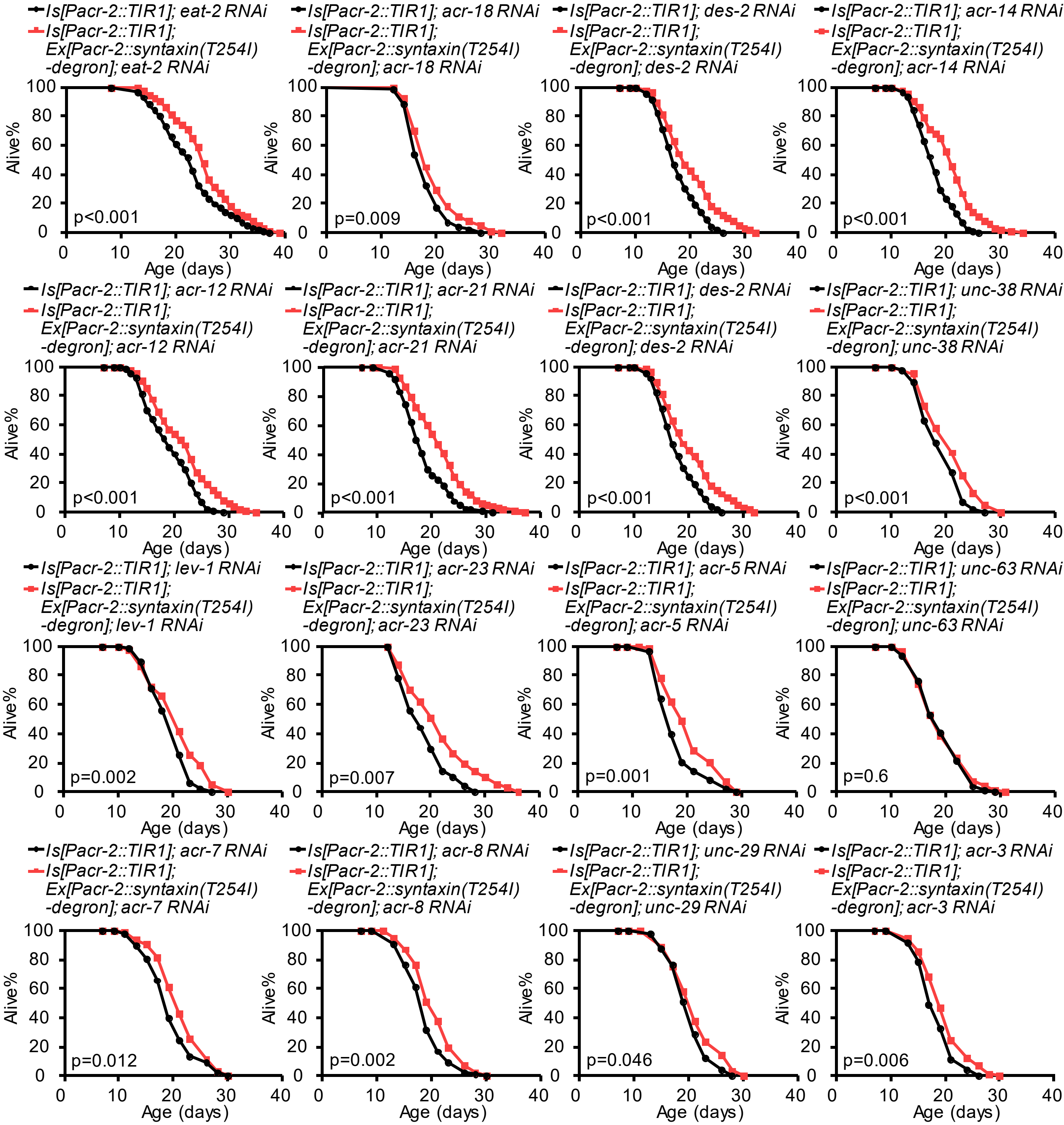


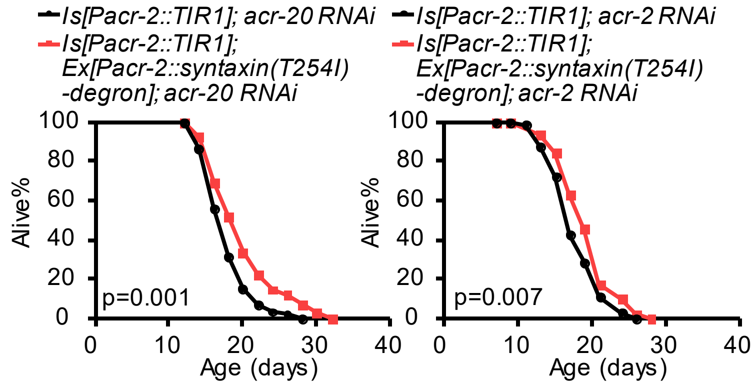


**Figure S6. RNAi screen of AChR genes that can block the lifespan-extension effect of cholinergic motor neurons.**
The *syntaxin(T254I)* transgene was induced from Day 7 adulthood as described in Figure 1C. *unc-63* was not selected as a candidate gene due to the poor growth of RNAi bacteria. For feeding RNAi experiments, NGM plates included carbenicillin (25 μg/ml) and IPTG (0.5 mM). HT115 bacteria-carrying vector or RNAi plasmid were seeded on RNAi plates 2 days before the experiment. Worms were fed RNAi bacteria from the egg stage. All lifespan assays were performed at 20°C. Logrank (Kaplan-Meier) was used to calculate p values.

Figure S7

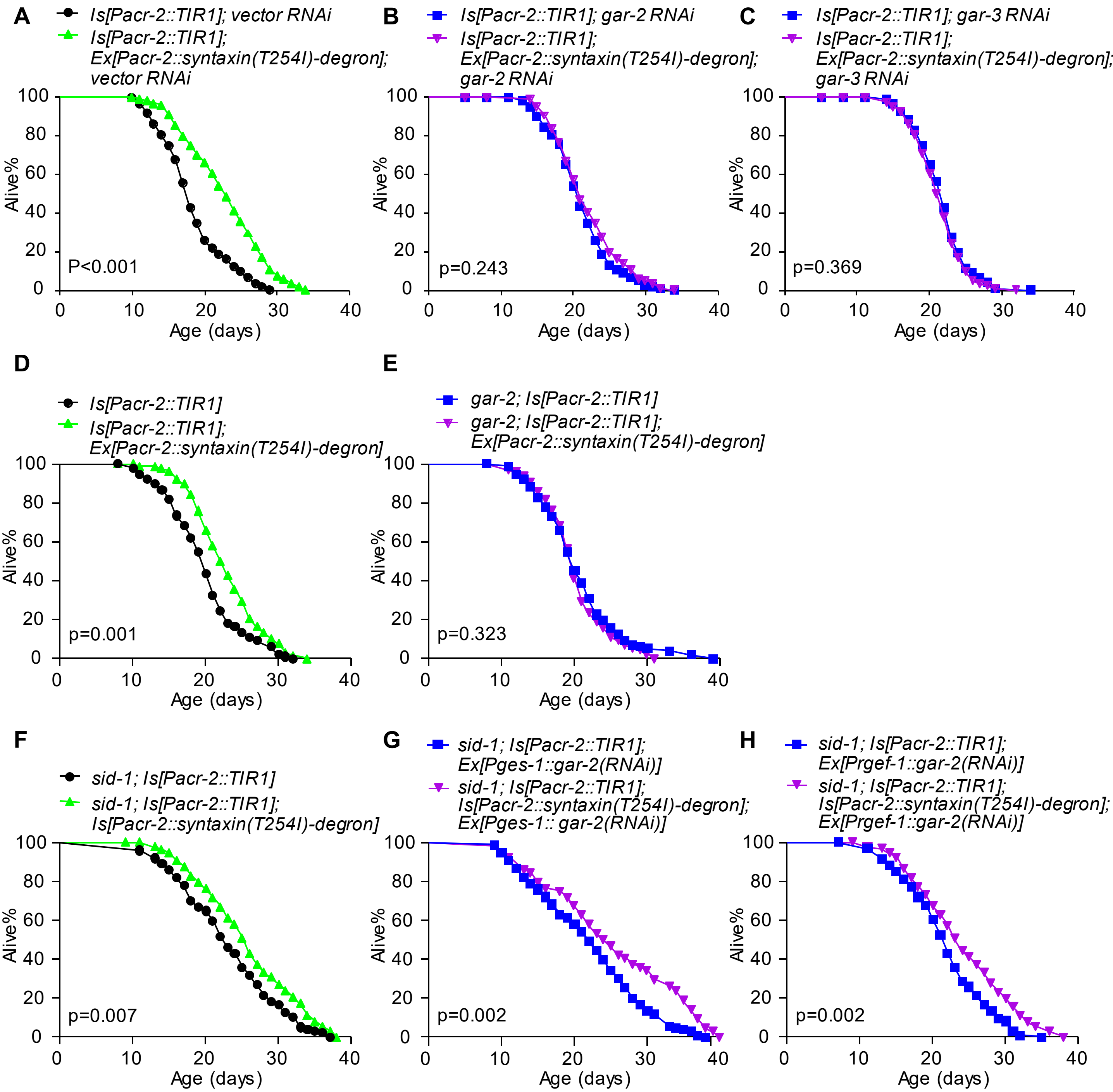


**Figure S7. Intestinal mAChR GAR-2 does not mediate the lifespan-extending effect of cholinergic motor neurons.**

(A) Enhancing the output of cholinergic motor neurons in mid-late life extends lifespan. The *syntaxin(T254I)* transgene was induced from D7 adulthood as described in Figure. 1C. (B-C) RNAi of *gar-2* (B) or *gar-3* (C) blocks the lifespan-extension effect of cholinergic motor neurons in mid-late life. For feeding RNAi experiments, NGM plates included carbenicillin (25 μg/ml) and IPTG (0.5 mM). HT115 bacteria-carrying vector or RNAi plasmid were seeded on RNAi plates 2 days before the experiment. Worms were fed RNAi bacteria from the egg stage. (D-E) Mutation in *gar-2* blocks the lifespan-extension effect of cholinergic motor neurons in mid-late life. (F-H) RNAi of *gar-2* neither in the intestine (G), nor in neurons (H), abolishes the lifespan-extension effect of cholinergic motor neurons in mid-late life (F). dsRNA against *gar-3* was expressed as a transgene specifically in the intestine and neurons using *ges-1* and *rgef-1* promoter, respectively. These experiments were carried out in a *sid-1* mutant background where systemic effect of RNAi is absent. All lifespan assays were performed at 20°C and were repeated at least twice. Log-rank (Kaplan-Meier) was used to calculate p values.

Figure S8

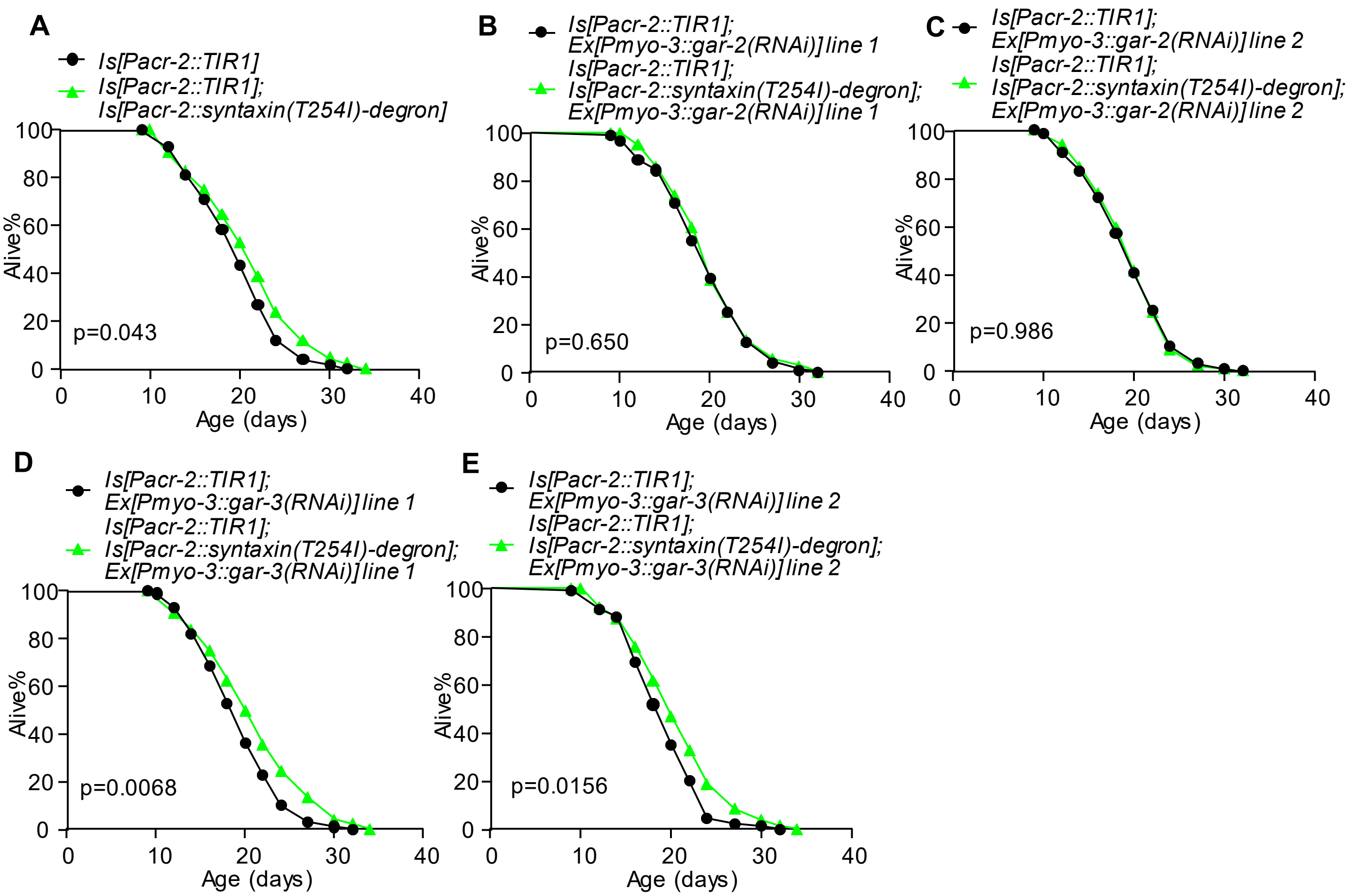


**Figure S8. Muscle mAChR GAR-2 mediate the lifespan-extending effect of cholinergic motor neurons, while muscle GAR-3 not.**

(A) Enhancing the output of cholinergic motor neurons in mid-late life extends lifespan. The *syntaxin(T254I)* transgene was induced from D7 adulthood as described in Figure 1C. (B-C) RNAi of *gar-2* in muscle blocks the lifespan-extension effect of cholinergic motor neurons in mid-late life. (D-E) RNAi of *gar-3* does not abolish the lifespan-extension effect of cholinergic motor neurons in mid-late life. dsRNA against *gar-2* or *gar-3* was expressed as a transgene specifically in muscle using *myo-3* promoter, respectively. All lifespan assays were performed at 20°C. Log-rank (Kaplan-Meier) was used to calculate p values.

Figure S9

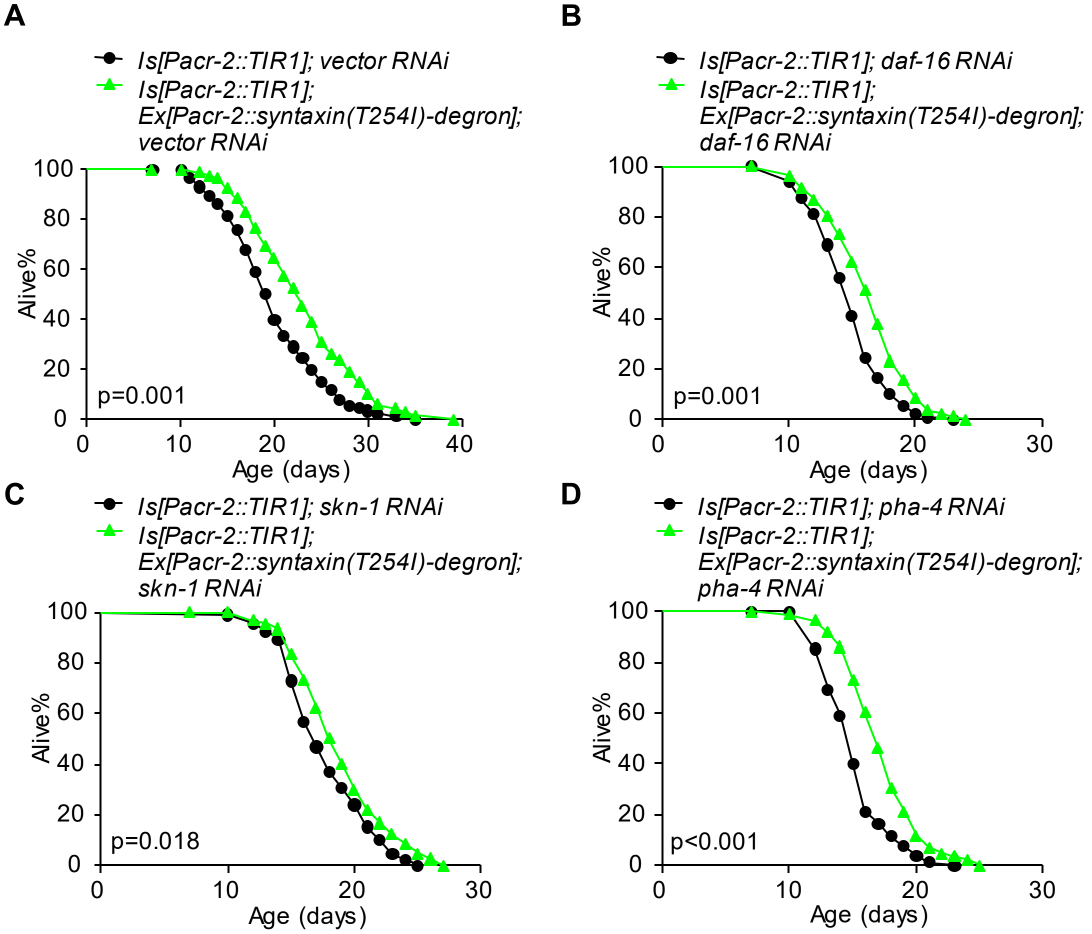


**Figure S9. The transcription factors DAF-16, SKN-1 and PHA-4 are not required for cholinergic motor neurons to promote lifespan in mid-late life.**
(A-D) Enhancing the output of cholinergic motor neurons in mid-late life extends lifespan (A). This effect cannot be suppressed by RNAi of *daf-16* (B), *skn-1* (C) or *pha-4* (D). The *syntaxin(T254I)* transgene was induced from D7 adulthood as described in Figure. 1C. For feeding RNAi experiments, NGM plates included carbenicillin (25 μg/ml) and IPTG (0.5 mM). HT115 bacteria-carrying vector or RNAi plasmid were seeded on RNAi plates 2 days before the experiment. Worms were fed RNAi bacteria from the L4 stage. All lifespan assays were performed at 20°C and were repeated at least twice. Logrank (Kaplan-Meier) was used to calculate p values.

Figure S10

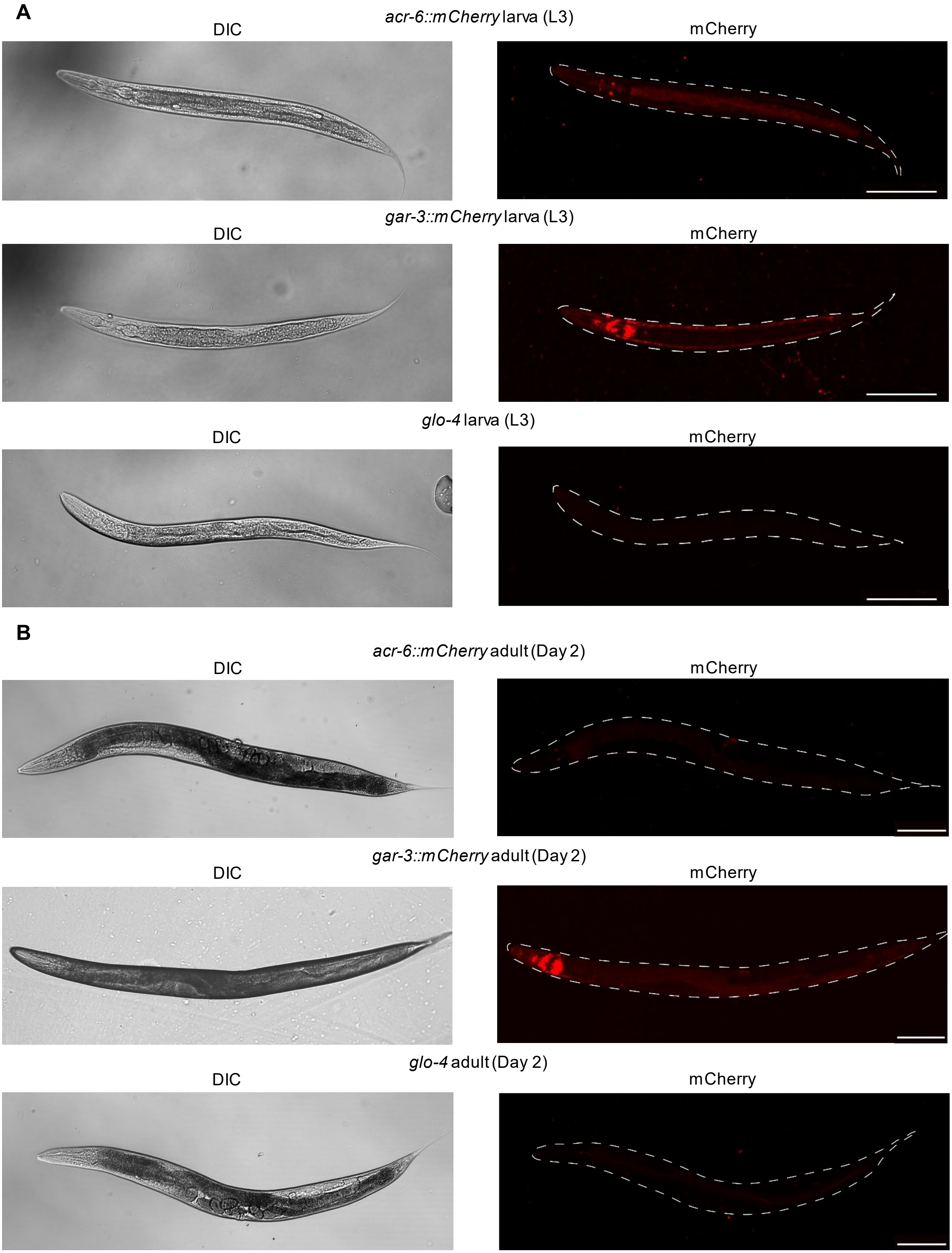


**Figure S10. The representative whole worm images of endogenous ACR-6 and GAR-3 proteins at L3 Larva stage (A) and Day 2 adult stage (B).**
*glo-4(ok623)* mutant worms were used as background with reduced gut auto-fluorescence. Scale bars, 100 μm.
