## Supplemental Table 1 for "Temporally controlled nervous system-to-gut signaling bidirectionally regulates longevity in *C. elegans*"

### Supplementary Table 1 Summary of Lifespan Results

| gene name | Mean Lifespan ±SEM | 25% Lifespan | Median Lifespan | 75% Lifespan | N (assayed /total) | P value | Figure |
|---|---|---|---|---|---|---|---|
| WT | 24.972±0.888 | 18 | 26 | 31 | 82/120 | | 1A |
| *Ex[Pacr-2::Tetx]* | 25.711±0.583 | 23 | 27 | 29 | 91/112 | 0.285 | 1A |
| WT | 20.184±0.931 | 14 | 19 | 26 | 70/100 | | Repeat 1A |
| *Ex[Pacr-2::Tetx]* | 22.928±0.699 | 20 | 23 | 28 | 61/80 | 0.144 | Repeat 1A |
| WT | 18.097±0.501 | 14 | 18 | 21 | 91/120 | | 1B |
| *Ex[Pacr-2::syntaxin(T254I)]* | 15.130±0.441 | 12 | 14 | 17 | 79/120 | <0.001 | 1B |
| WT | 21.599±0.414 | 19 | 22 | 24 | 76/100 | | Repeat 1B |
| *Ex[Pacr-2::syntaxin(T254I)]* | 18.294±0.568 | 15 | 18 | 22 | 75/100 | 0.002 | Repeat 1B |
| *Is[Pacr-2::TIR1]* | 23.821±0.716 | 19 | 24 | 28 | 93/110 | | 1D |
| *Is[Pacr-2::TIR1]; Ex[Pacr-2::syntaxin(T254I)-degron]* | 23.270±0.712 | 19 | 24 | 27 | 83/105 | 0.462 | 1D |
| *Is[Pacr-2::TIR1]* | 19.968±0.673 | 15 | 20 | 23 | 73/100 | | Repeat 1D |
| *Is[Pacr-2::TIR1]; Ex[Pacr-2::syntaxin(T254I)-degron]* | 19.670±0.612 | 16 | 19 | 23 | 67/100 | 0.559 | Repeat 1D |
| *Is[Pacr-2::TIR1]* | 21.524±0.615 | 17 | 22 | 26 | 86/120 | | 1E |
| *Is[Pacr-2::TIR1]; Ex[Pacr-2::syntaxin(T254I)-degron]* | 18.270±0.719 | 14 | 17 | 21 | 57/100 | 0.001 | 1E |
| *Is[Pacr-2::TIR1]* | 24.050±0.813 | 19 | 25 | 28 | 67/120 | | Repeat 1E |
| *Is[Pacr-2::TIR1]; Ex[Pacr-2::syntaxin(T254I)-degron]* | 19.944±0.476 | 17 | 20 | 23 | 72/120 | <0.001 | Repeat 1E |
| *Is[Pacr-2::TIR1]* | 21.959±0.897 | 16 | 23 | 26 | 42/100 | | 1F |
| *Is[Pacr-2::TIR1]; Ex[Pacr-2::syntaxin(T254I)-degron]* | 18.712±0.805 | 14 | 18 | 22 | 50/100 | 0.013 | 1F |

| | | | | | | | |
|---|---|---|---|---|---|---|---|
| *Is[Pacr-2::TIR1]* | 21.304±0.631 | 18 | 22 | 25 | 66/100 | | Repeat 1F |
| *Is[Pacr-2::TIR1]; Ex[Pacr-2::syntaxin(T254I)-degron]* | 19.689±0.761 | 15 | 19 | 24 | 57/100 | 0.256 | Repeat 1F |
| *Is[Pacr-2::TIR1]* | 20.212±0.771 | 16 | 20 | 24 | 66/110 | | 1G |
| *Is[Pacr-2::TIR1]; Ex[Pacr-2::syntaxin(T254I)-degron]* | 18.578±0.650 | 15 | 18 | 22 | 48/110 | 0.074 | 1G |
| *Is[Pacr-2::TIR1]* | 18.866±0.669 | 15 | 18 | 21 | 62/90 | | Repeat 1G |
| *Is[Pacr-2::TIR1]; Ex[Pacr-2::syntaxin(T254I)-degron]* | 19.461±0.655 | 16 | 19 | 23 | 66/120 | 0.526 | Repeat 1G |
| *Is[Pacr-2::TIR1]* | 23.487±0.590 | 19 | 24 | 27 | 108/120 | | 1H |
| *Is[Pacr-2::TIR1]; Ex[Pacr-2::syntaxin(T254I)-degron]* | 22.900±0.746 | 17 | 21 | 28 | 110/120 | 0.746 | 1H |
| *Is[Pacr-2::TIR1]* | 22.246±0.732 | 17 | 22 | 27 | 82/120 | | Repeat 1H |
| *Is[Pacr-2::TIR1]; Ex[Pacr-2::syntaxin(T254I)-degron]* | 21.665±0.964 | 15 | 19 | 27 | 77/110 | 0.663 | Repeat 1H |
| *Is[Pacr-2::TIR1]* | 23.769±0.580 | 20 | 24 | 27 | 104/120 | | 1I |
| *Is[Pacr-2::TIR1]; Ex[Pacr-2::syntaxin(T254I)-degron]* | 22.737±0.566 | 18 | 23 | 26 | 98/120 | 0.210 | 1I |
| *Is[Pacr-2::TIR1]* | 21.115±0.695 | 17 | 21 | 25 | 76/100 | | Repeat 1I |
| *Is[Pacr-2::TIR1]; Ex[Pacr-2::syntaxin(T254I)-degron]* | 22.044±0.550 | 18 | 22 | 25 | 84/120 | 0.702 | Repeat 1I |
| *Is[Pacr-2::TIR1]* | 20.174±0.668 | 16 | *20* | 23 | 61/120 | | 1J |
| *Is[Pacr-2::TIR1]; Ex[Pacr-2::syntaxin(T254I)-degron]* | 23.370±0.849 | 20 | *23* | 27 | 65/120 | 0.002 | 1J |
| *Is[Pacr-2::TIR1]* | 22.275±0.604 | 17 | 22 | 27 | 95/100 | | Repeat 1J |

| | | | | | | | |
|---|---|---|---|---|---|---|---|
| *Is[Pacr-2::TIR1]; Ex[Pacr-2::syntaxin(T254I)-degron]* | 24.262±0.656 | 19 | 24 | 29 | 93/105 | 0.021 | Repeat 1J |
| *Is[Pacr-2::TIR1]* | 19.616±0.528 | 16 | 21 | 22 | 65/75 | | 1K |
| *Is[Pacr-2::TIR1]; Ex[Pacr-2::syntaxin(T254I)-degron]* | 22.588±0.555 | 20 | 22 | 24 | 47/60 | 0.002 | 1K |
| *Is[Pacr-2::TIR1]* | 21.561±0.652 | 16 | 22 | 27 | 96/105 | | Repeat 1K |
| *Is[Pacr-2::TIR1]; Ex[Pacr-2::syntaxin(T254I)-degron]* | 25.063±0.642 | 21 | 24 | 29 | 96/110 | 0.001 | Repeat 1K |
| *Is[Pacr-2::TIR1]* | 21.384±0.700 | 21 | 21 | 24 | 53/60 | | 1L |
| *Is[Pacr-2::TIR1]; Ex[Pacr-2::syntaxin(T254I)-degron]* | 23.519±0.527 | 18 | 23 | 26 | 62/75 | 0.066 | 1L |
| *Is[Pacr-2::TIR1]* | 23.568±0.611 | 19 | 24 | 28 | 79/100 | | Repeat 1L |
| *Is[Pacr-2::TIR1]; Ex[Pacr-2::syntaxin(T254I)-degron]* | 24.987±0.611 | 20 | 25 | 29 | 86/105 | 0.084 | Repeat 1L |
| WT | 21.528±0.529 | 18 | 21 | 25 | 87/120 | | 2A |
| *Ex[Pacr-2::syntaxin(T254I)]* | 18.279±0.480 | 15 | 18 | 22 | 92/120 | <0.001 | 2A |
| WT | 20.560±0.555 | 16 | 21 | 24 | 82/102 | | Repeat 2A |
| *Ex[Pacr-2::syntaxin(T254I)]* | 18.532±0.506 | 14 | 18 | 22 | 94/100 | 0.013 | Repeat 2A |
| *unc-31* | 25.966±1.076 | 16 | 27 | 34 | 81/100 | | 2B |
| *unc-31; Ex[Pacr-2::syntaxin(T254I)]* | 21.215±0.872 | 14 | 21 | 27 | 79/80 | <0.001 | 2B |
| *unc-31* | 25.848±1.337 | 17 | 23 | 36 | 68/80 | | Repeat 2B |
| *unc-31; Ex[Pacr-2::syntaxin(T254I)]* | 19.548±0.896 | 13 | 16 | 25 | 95/100 | <0.001 | Repeat 2B |
| *unc-17* | 20.458±0.552 | 16 | 20 | 24 | 65/120 | | 2C |
| *unc-17; Ex[Pacr-2::syntaxin(T254I)]* | 20.476±0.502 | 16 | 20 | 24 | 78/120 | 0.966 | 2C |
| *unc-17* | 19.055±0.550 | 15 | 18 | 23 | 88/100 | | Repeat 2C |

| | | | | | | | |
|---|---|---|---|---|---|---|---|
| *unc-17; Ex[Pacr-2::syntaxin(T254I)]* | 19.261±0.523 | 15 | 19 | 23 | 86/95 | 0.914 | Repeat 2C |
| *acr-6* | 19.235±0.727 | 15 | 19 | 23 | 59/120 | | 2D |
| *acr-6*; *Ex[Pacr-2::syntaxin(T254I)]* | 19.389±0.750 | 14 | 19 | 23 | 78/120 | 0.632 | 2D |
| *acr-6* | 20.050±0.813 | 14 | 17 | 26 | 57/90 | | Repeat 2D |
| *acr-6*; *Ex[Pacr-2::syntaxin(T254I)]* | 18.774±0.795 | 13 | 17 | 23 | 64/100 | 0.238 | Repeat 2D |
| *acr-6; Ex[Pges-1::acr-6]* | 23.178±0.982 | 18 | 24 | 29 | 52/120 | | 2E |
| *acr-6; Ex[Pacr-2:: syntaxin(T254I)]; Ex[Pges-1::acr-6]* | 19.521±0.794 | 14 | 19 | 24 | 70//120 | 0.007 | 2E |
| *acr-6; Ex[Pges-1::acr-6]* | 20.813±0.983 | 13 | 20 | 28 | 61/90 | | Repeat 2E |
| acr-6; *Ex[Pacr-2:: syntaxin(T254I)]*; *Ex[Pges-1::acr-6]* | 16.996±0.718 | 12 | 16 | 21 | 70/120 | 0.002 | Repeat 2E |
| *acr-6; Ex[Prgef-1::acr-6]* | 18.530±0.998 | 12 | 18 | 24 | 52/120 | | 2F |
| acr-6; *Ex[Pacr-2:: syntaxin(T254I)]*; *Ex[Prgef-1::acr-6]* | 19.455±0.963 | 14 | 19 | 23 | 43/120 | 0.680 | 2F |
| *acr-6; Ex[Prgef-1::acr-6]* | 17.152±0.937 | 12 | 14 | 23 | 53/90 | | Repeat 2F |
| acr-6; *Ex[Pacr-2:: syntaxin(T254I)]*; *Ex[Prgef-1::acr-6]* | 15.941±0.958 | 11 | 13 | 21 | 43/75 | 0.419 | Repeat 2F |
| *vector RNAi* | 21.838±0.521 | 18 | 21 | 25 | 74/120 | | 3A |
| *Ex[Pacr-2:: syntaxin(T254I)]; vector RNAi* | 17.789±0.551 | 15 | 17 | 21 | 71/105 | <0.001 | 3A |
| *vector RNAi* | 21.320±0.478 | 18 | 21 | 25 | 75/100 | | Repeat 3A |
| *Ex[Pacr-2:: syntaxin(T254I)]; vector RNAi* | 16.473±0.443 | 14 | 15 | 19 | 78/100 | <0.001 | Repeat 3A |
| *daf-16 RNAi* | 16.703±0.367 | 13 | 17 | 20 | 104/120 | | 3B |
| *Ex[Pacr-2:: syntaxin(T254I)]; daf-16 RNAi* | 17.278±0.388 | 14 | 17 | 21 | 91/120 | 0.260 | 3B |
| *daf-16 RNAi* | 14.287±0.257 | 12 | 14 | 16 | 108/120 | | Repeat 3B |

| | | | | | | | |
|---|---|---|---|---|---|---|---|
| *Ex[Pacr-2:: syntaxin(T254I)]; daf-16 RNAi* | 14.598±0.284 | 12 | 14 | 17 | 107/118 | 0.291 | Repeat 3B |
| *hsf-1 RNAi* | 14.329±0.247 | 13 | 14 | 16 | 79/120 | | 3C |
| *Ex[Pacr-2:: syntaxin(T254I)]; hsf-1 RNAi* | 13.053±0.235 | 11 | 13 | 15 | 76/120 | <0.001 | 3C |
| *hsf-1 RNAi* | 14.550±0.250 | 13 | 14 | 16 | 108/120 | | Repeat 3C |
| *Ex[Pacr-2:: syntaxin(T254I)]; hsf-1 RNAi* | 13.426±0.227 | 12 | 13 | 15 | 99/120 | 0.001 | Repeat 3C |
| *skn-1 RNAi* | 18.461±0.341 | 16 | 18 | 21 | 115/120 | | 3D |
| *Ex[Pacr-2:: syntaxin(T254I)]; skn-1 RNAi* | 16.849±0.382 | 14 | 16 | 19 | 86/120 | 0.006 | 3D |
| *skn-1 RNAi* | 20.402±0.465 | 17 | 19 | 23 | 87/120 | | Repeat 3D |
| *Ex[Pacr-2:: syntaxin(T254I)]; skn-1 RNAi* | 16.200±0.277 | 15 | 17 | 17 | 102/120 | <0.001 | Repeat 3D |
| *pha-4 RNAi* | 19.083±0.414 | 15 | 19 | 22 | 108/120 | | 3E |
| *Ex[Pacr-2:: syntaxin(T254I)]; pha-4 RNAi* | 17.330±0.376 | 14 | 17 | 20 | 97/120 | 0.002 | 3E |
| *pha-4 RNAi* | 24.891±0.642 | 21 | 25 | 29 | 92/120 | | Repeat 3E |
| *Ex[Pacr-2:: syntaxin(T254I)]; pha-4 RNAi* | 20.696±0.497 | 17 | 21 | 23 | 92/120 | <0.001 | Repeat 3E |
| *sid-1* | 20.692±0.821 | 15 | 19 | 27 | 85/100 | | 3G |
| *sid-1; Ex[Pacr-2::Tetx]* | 21.948±0.597 | 17 | 21 | 27 | 97/100 | 0.876 | 3G |
| *sid-1* | 22.409±0.646 | 17 | 23 | 27 | 90/107 | | Repeat 3G |
| *sid-1; Ex[Pacr-2::Tetx]* | 22.977±0.486 | 21 | 23 | 27 | 87/100 | 0.586 | Repeat 3G |
| *sid-1; Ex[Pvha-6::daf-16(RNAi)]* | 16.833±0.398 | 13 | 17 | 19 | 96/100 | | 3H |
| *sid-1; Ex[Pacr-2::Tetx]; Ex[Pvha-6::daf-16(RNAi)]* | 16.140±0.386 | 13 | 15 | 19 | 100/100 | 0.245 | 3H |
| *sid-1; Ex[Pvha-6::daf-16(RNAi)]* | 16.940±0.401 | 15 | 17 | 19 | 100/100 | | Repeat 3H |

| | | | | | | | |
|---|---|---|---|---|---|---|---|
| *sid-1; Ex[Pacr-2::Tetx]; Ex[Pvha-6::daf-16(RNAi)]* | 16.400±0.382 | 13 | 17 | 19 | 100/100 | 0.288 | Repeat 3H |
| *Is[Pacr-2::TIR1]* | 22.722±0.639 | 18 | 23 | 28 | 92/100 | | 4A |
| *Is[Pacr-2::TIR1]; Is[Pacr-2:: syntaxin(T254I)-degron]* | 25.805±0.753 | 22 | 26 | 30 | 70/81 | 0.001 | 4A |
| *Is[Pacr-2::TIR1]* | 20.667±0.660 | 15 | 20 | 27 | 105/120 | | Repeat 4A |
| *Is[Pacr-2::TIR1]; Is[Pacr-2:: syntaxin(T254I)-degron]* | 23.545±0.808 | 18 | 23 | 28 | 89/120 | 0.014 | Repeat 4A |
| *unc-17; Is[Pacr-2::TIR1]* | 20.215±0.639 | 15 | 21 | 25 | 93/100 | | 4B |
| *unc-17; Is[Pacr-2::TIR1]; Is[Pacr-2:: syntaxin(T254I)-degron]* | 20.813±0.725 | 14 | 21 | 27 | 86/90 | 0.258 | 4B |
| *unc-17; Is[Pacr-2::TIR1]* | 21.491±0.576 | 17 | 22 | 26 | 107/120 | | Repeat 4B |
| *unc-17; Is[Pacr-2::TIR1]; Is[Pacr-2:: syntaxin(T254I)-degron]* | 20.813±0.535 | 17 | 21 | 25 | 102/120 | 0.187 | Repeat 4B |
| *gar-3; Is[Pacr-2::TIR1]* | 21.170±0.436 | 18 | 21 | 25 | 94/105 | | 4C |
| *gar-3; Is[Pacr-2::TIR1]; Ex[Pacr-2:: syntaxin(T254I)-degron]* | 21.207±0.621 | 17 | 21 | 25 | 70/90 | 0.535 | 4C |
| *gar-3; Is[Pacr-2::TIR1]* | 21.247±0.649 | 16 | 21 | 26 | 80/95 | | Repeat 4C |
| *gar-3; Is[Pacr-2::TIR1]; Ex[Pacr-2:: syntaxin(T254I)-degron]* | 21.326±0.595 | 16 | 22 | 25 | 89/95 | 0.904 | Repeat 4C |
| *sid-1; Is[Pacr-2::TIR1]* | 23.110±0.665 | 18 | 23 | 28 | 98/100 | | 4D |
| *sid-1; Is[Pacr-2::TIR1]; Is[Pacr-2:: syntaxin(T254I)-degron]* | 25.791±0.705 | 21 | 26 | 31 | 88/101 | 0.007 | 4D |
| *sid-1; Is[Pacr-2::TIR1]* | 21.476±0.661 | 16 | 21 | 27 | 107/120 | | Repeat 4D |
| *sid-1; Is[Pacr-2::TIR1]; Is[Pacr-2:: syntaxin(T254I)-degron]* | 24.590±0.763 | 19 | 24 | 30 | 92/105 | 0.003 | Repeat 4D |
| *sid-1; Is[Pacr-2::TIR1]; Ex[Pges-1::gar-3(RNAi)]* | 23.507±0.920 | 17 | 24 | 30 | 82/86 | | 4E |
| *sid-1; Is[Pacr-2::TIR1]; Is[Pacr-2::* | 23.604±1.223 | 16 | 24 | 30 | 47/60 | 0.917 | 4E |

| | | | | | | | |
|---|---|---|---|---|---|---|---|
| *syntaxin(T254I)-degron]; Ex[Pges-1::gar-3(RNAi)]* | | | | | | | |
| *sid-1; Is[Pacr-2::TIR1]; Ex[Pges-1::gar-3(RNAi)]* | 20.596±0.704 | 17 | 20 | 23 | 64/100 | | Repeat 4E |
| *sid-1; Is[Pacr-2::TIR1]; Is[Pacr-2:: syntaxin(T254I)-degron]; Ex[Pges-1::gar-3(RNAi)]* | 20.939±1.009 | 16 | 19 | 24 | 48/90 | 0.754 | Repeat 4E |
| *sid-1; Is[Pacr-2::TIR1]; Ex[Prgef-1::gar-3(RNAi)]* | 22.000±0.526 | 19 | 21 | 24 | 103/103 | | 4F |
| *sid-1; Is[Pacr-2::TIR1]; Is[Pacr-2:: syntaxin(T254I)-degron]; Ex[Prgef-1::gar-3(RNAi)]* | 24.926±0.578 | 22 | 25 | 29 | 101/107 | <0.001 | 4F |
| *sid-1; Is[Pacr-2::TIR1]; Ex[Prgef-1::gar-3(RNAi)]* | 18.669±0.583 | 14 | 17 | 23 | 93/101 | | Repeat 4F |
| *sid-1; Is[Pacr-2::TIR1]; Is[Pacr-2:: syntaxin(T254I)-degron]; Ex[Prgef-1::gar-3(RNAi)]* | 21.813±0.584 | 17 | 23 | 26 | 86/98 | 0.002 | Repeat 4F |
| *Is[Pacr-2::TIR1]; vector RNAi* | 20.012±0.557 | 17 | 19 | 23 | 86/96 | | 5A |
| *Is[Pacr-2::TIR1]; Ex[Pacr-2:: syntaxin(T254I)-degron]; vector RNAi* | 23.110±0.655 | 19 | 23 | 27 | 73/94 | 0.001 | 5A |
| *Is[Pacr-2::TIR1]; vector RNAi* | 20.182±0.511 | 16 | 19 | 23 | 97/120 | | Repeat 5A |
| *Is[Pacr-2::TIR1]; Ex[Pacr-2:: syntaxin(T254I)-degron]; vector RNAii* | 24.045±0.616 | 20 | 23 | 28 | 93/120 | <0.001 | Repeat 5A |
| *Is[Pacr-2::TIR1]; hsf-1 RNAi* | 12.982±0.163 | 12 | 13 | 14 | 81/95 | | 5B |
| *Is[Pacr-2::TIR1]; Ex[Pacr-2:: syntaxin(T254I)-degron]; hsf-1 RNAi* | 13.004±0.156 | 12 | 13 | 14 | 87/96 | 0.997 | 5B |
| *Is[Pacr-2::TIR1]; hsf-1 RNAi* | 15.688±0.303 | 14 | 16 | 18 | 93/105 | | Repeat 5B |
| *Is[Pacr-2::TIR1]; Ex[Pacr-2::* | 16.027±0.305 | 14 | 16 | 18 | 81/105 | 0.564 | Repeat 5B |

| | | | | | | | |
|---|---|---|---|---|---|---|---|
| *syntaxin(T254I)-degron]; hsf-1 RNAi* | | | | | | | |
| *sid-1; Is[Pacr-2::TIR1]* | 16.046±0.614 | 12 | 14 | 18 | 91/100 | | 5D |
| *sid-1; Is[Pacr-2::TIR1]; Is[Pacr-2:: syntaxin(T254I)-degron]* | 21.380±0.635 | 16 | 20 | 26 | 100/102 | <0.001 | 5D |
| *sid-1; Is[Pacr-2::TIR1]* | 16.860±0.632 | 12 | 16 | 20 | 86/100 | | Repeat 5D |
| *sid-1; Is[Pacr-2::TIR1]; Is[Pacr-2:: syntaxin(T254I)-degron]* | 19.510±0.624 | 14 | 18 | 24 | 98/100 | 0.007 | Repeat 5D |
| *sid-1; Is[Pacr-2::TIR1]; Ex[Pges-1::hsf-1(RNAi)]* | 14.353±0.405 | 12 | 14 | 16 | 85/100 | | 5E |
| *sid-1; Is[Pacr-2::TIR1]; Is[Pacr-2:: syntaxin(T254I)-degron]; Ex[Pges-1::hsf-1(RNAi)]* | 14.545±0.411 | 12 | 14 | 18 | 84/100 | 0.878 | 5E |
| *sid-1; Is[Pacr-2::TIR1]; Ex[Pges-1::hsf-1(RNAi)]* | 14.634±0.381 | 12 | 14 | 16 | 60/100 | | Repeat 5E |
| *sid-1; Is[Pacr-2::TIR1]; Is[Pacr-2:: syntaxin(T254I)-degron]; Ex[Pges-1::hsf-1(RNAi)]* | 14.724±0.402 | 12 | 14 | 16 | 77/100 | 0.813 | Repeat 5E |
| *sid-1; Is[Pacr-2::TIR1]; Ex[Prgef-1::hsf-1(RNAi)]* | 20.231±0.536 | 16 | 20 | 24 | 91/100 | | 5F |
| *sid-1; Is[Pacr-2::TIR1]; Is[Pacr-2:: syntaxin(T254I)-degron]; Ex[Prgef-1::hsf-1(RNAi)]* | 24.782±0.563 | 21 | 25 | 29 | 87/100 | <0.001 | 5F |
| *sid-1; Is[Pacr-2::TIR1]; Ex[Prgef-1::hsf-1(RNAi)]* | 20.139±0.671 | 14 | 19 | 26 | 100/103 | | Repeat 5F |
| *sid-1; Is[Pacr-2::TIR1]; Is[Pacr-2:: syntaxin(T254I)-degron]; Ex[Prgef-1::hsf-1(RNAi)]* | 22.705±0.745 | 17 | 23 | 27 | 93/101 | 0.014 | Repeat 5F |

The Log Rank (Mantel-Cox) test was used for statistical analysis. N numbers are described as: assayed/total, i.e. the number of assayed animals/total animals included to the plates initially. The difference represents the number of animals that were censored (crawled off the plate, bagged or exploded). Lifespan data was analyzed with GraphPad Prism 8 (GraphPad Software, Inc.) and IBM SPSS Statistics 21 (IBM, Inc.). Log-rank (Kaplan-Meier) was used to calculate *P* values. 5-Fluoro-2'-deoxyuridine (FUDR) was included in assays involving TeTx transgene worms, *unc-31* and *unc-17* mutant worms, which show a defect in egg laying.
